# *Snrpb*, the CCMS gene, is required in neural crest cells for proper splicing of genes essential for craniofacial morphogenesis

**DOI:** 10.1101/2021.08.07.455514

**Authors:** Sabrina Shameen Alam, Shruti Kumar, Marie-Claude Beauchamp, Eric Bareke, Alexia Boucher, Nadine Nzirorera, Reinnier Padilla, Si Jing Zhang, Jacek Majewski, Loydie A. Jerome-Majewska

## Abstract

Heterozygous mutations in *SNRPB*, an essential core component of the five small ribonucleoprotein particles of the spliceosome, are responsible for Cerebrocostomandibular Syndrome (CCMS). We show that *Snrpb* heterozygous embryos arrest shortly after implantation. Additionally, heterozygous deletion of *Snrpb* in the developing brain and neural crest cells models craniofacial malformations found in CCMS, and results in death shortly after birth. RNAseq analysis of mutant heads prior to morphological defects revealed increased exon-skipping and intron-retention in association with increased 5’ splice site strength. We found increased exon-skipping in negative regulators of the P53-pathway, increased levels of nuclear P53, P53-target genes. However, removing *TrP53* in *Snrpb* heterozygous mutant neural crest cells did not completely rescue craniofacial development. We also found a small but significant increase in exon-skipping of several transcripts required for head and midface development, including *Smad2* and *Rere*. Furthermore, mutant embryos exhibited ectopic or missing expression of *Fgf8* and *Shh*, which are required to coordinate face and brain development. Thus, we propose that mis-splicing of transcripts that regulate P53-activity and craniofacial-specific genes both contribute to craniofacial malformations.

## INTRODUCTION

95% of human pre-mRNAs are alternatively spliced to generate multiple mRNAs (Chen and Manley, 2009; Nilsen and Graveley, 2010) thus increasing the number and diversity of proteins expressed from the human genome. The major spliceosome or U2-dependent spliceosome catalyses 99% of RNA splicing reactions in human (Wickramasinghe *et al*., 2015), while the minor or U12-dependent spliceosome is responsible for splicing of approximately 700 minor introns in 666 genes (Olthof *et al*., 2019). The major spliceosome is composed of U1, U2, U5, and U4/U6 <u>s</u>mall <u>n</u>uclear <u>r</u>ibo<u>n</u>ucleo<u>p</u>rotein proteins (snRNPs) named for their core associated small RNAs (snRNAs). *SNRPB* encodes SmB and SmB’ which are core components of the spliceosome. SmB/B’ help to form the heptameric ring on the U snRNAs of the five snRNPs of the major spliceosome. Several groups have reported heterozygous mutations in *SNRPB* in patients with Cerebrocostomandibular Syndrome (CCMS, OMIM# 117650) (Lynch *et al*., 2014, Barcot *et al*., 2015, Tooley *et al*., 2016). CCMS patients have rib gaps and narrow chests, and craniofacial defects such as malar hypoplasia and micrognathia, with variable expressivity (Beauchamp *et al*., 2020) and incomplete penetrance.

In addition to the two coding transcripts, SmB and SmB’, *SNRPB* also encodes for a third premature termination codon (PTC) containing transcript that is predicted to undergo non-sense mediated decay. Most mutations found in CCMS patients increase inclusion of the PTC containing alternative exon 2 leading to no change or reduced levels of the coding transcripts in patient’s fibroblasts (Barcot *et al*., 2015 and Lynch *et al*. 2014). However, though it is presumed that increased expression of the PTC-containing transcript leads to reduced levels of SmB/SmB’ in all CCMS patients, reduced levels of SNRPB protein has not been reported in any CCMS patient cells. We postulated that a mouse model carrying mutation in *Snrpb* can be used to understand the role of *Snrpb* during embryogenesis and shed insight into the pathophysiology of CCMS.

Towards this goal, we generated a conditional mutant mouse line with *loxP* sequences flanking the genomic region that encompasses exons 2 and 3 of *Snrpb*. Using *β-Actin-cre* we showed that widespread heterozygous deletion (*Snrpb^+/-^*) of these exons reduced levels of *Snrpb* and resulted in embryonic arrest by embryonic day (E) 9.5. To investigate the role of *Snrpb* specifically during craniofacial development, we used *Wnt1-Cre2* to generate *Snrpb* heterozygosity in the developing brain and neural crest cells (*Snrpb^ncc+/-^*). We showed that a subset of these embryos survived to birth and died shortly after. Most *Snrpb^ncc+/-^* mutant embryos die between E17.5 and birth, with brain and craniofacial defects of variable expressivity. RNAseq analysis of the E9.0 embryonic heads of *Snrpb^ncc+/-^* mutants before morphological defects were apparent, revealed a significant increase in differential splicing events (DSE), but few differentially expressed genes (DEGs). Pathway analysis indicated that these DEGs were associated with the spliceosome and the P53-pathway. However, although nuclear P53 and apoptosis were increased in *Snrpb^ncc+/-^* embryos, reducing levels of P53 genetically in the neural crest did not rescue craniofacial defects in these mutants. Intriguingly, a number of DSE were found in genes important for craniofacial development. Furthermore, the expression of *Shh* and *Fgf8* which forms the facial organizer was disrupted in the craniofacial region. Our findings support disrupted splicing as the major driver of abnormalities in *Snrpb* mutant embryos. We suggest that abnormal splicing of genes important for craniofacial development results in an additive effect that disrupts morphogenesis of the head and brain.

## RESULTS

### Mouse embryos with deletion of exons 2 and 3 of *Snrpb* (*Snrpb^+/-^*) have reduced levels of *Snrpb* and die post-implantation

To test if reduced levels of *Snrpb* in mouse recapitulates abnormalities found in CCMS patients, we first generated a conditional mutant mouse line with *loxP* sequences in intron 1 and intron 3 of the gene. We then used *β-Actin*-cre to delete the *loxP*-flanked region – exon 2, the PTC encoding alternative exon 2, and exon 3 of *Snrpb* (Fig. 1A) – to produce *Snrpb* heterozygous mice (*Snrpb^+/-^*). However, no *Snrpb^+/-^* mice were found at birth (P0) or at weaning, indicating that these mutants died before birth (data not shown). To determine when *Snrpb^+/-^* embryos die, we dissected pregnant females from E6.5 – E10.5. We found that mutant *Snrpb^+/-^* embryos were undergoing resorption by E9.5 and were all dead by E10.5 (data not shown). We confirmed that *Cre*-mediated deletion of the *loxP*-flanked region generated a shorter *Snrpb* transcript of 527 bp (Fig. 1B) and resulted in a statistically significant 70% reduction in levels of *Snrpb* in E8.5 *Snrpb^+/-^* embryos (p= 0.0052, T-Test) (Fig. 1C). Thus, deletion of exons 2 – 3 of *Snrpb* leads to a significant reduction in *Snrpb* level in heterozygous mutant embryos. Our data indicate that the amount of functional protein expressed by a single wild-type allele of *Snrpb* was insufficient for embryonic growth and survival post-implantation.

**Figure 1:**
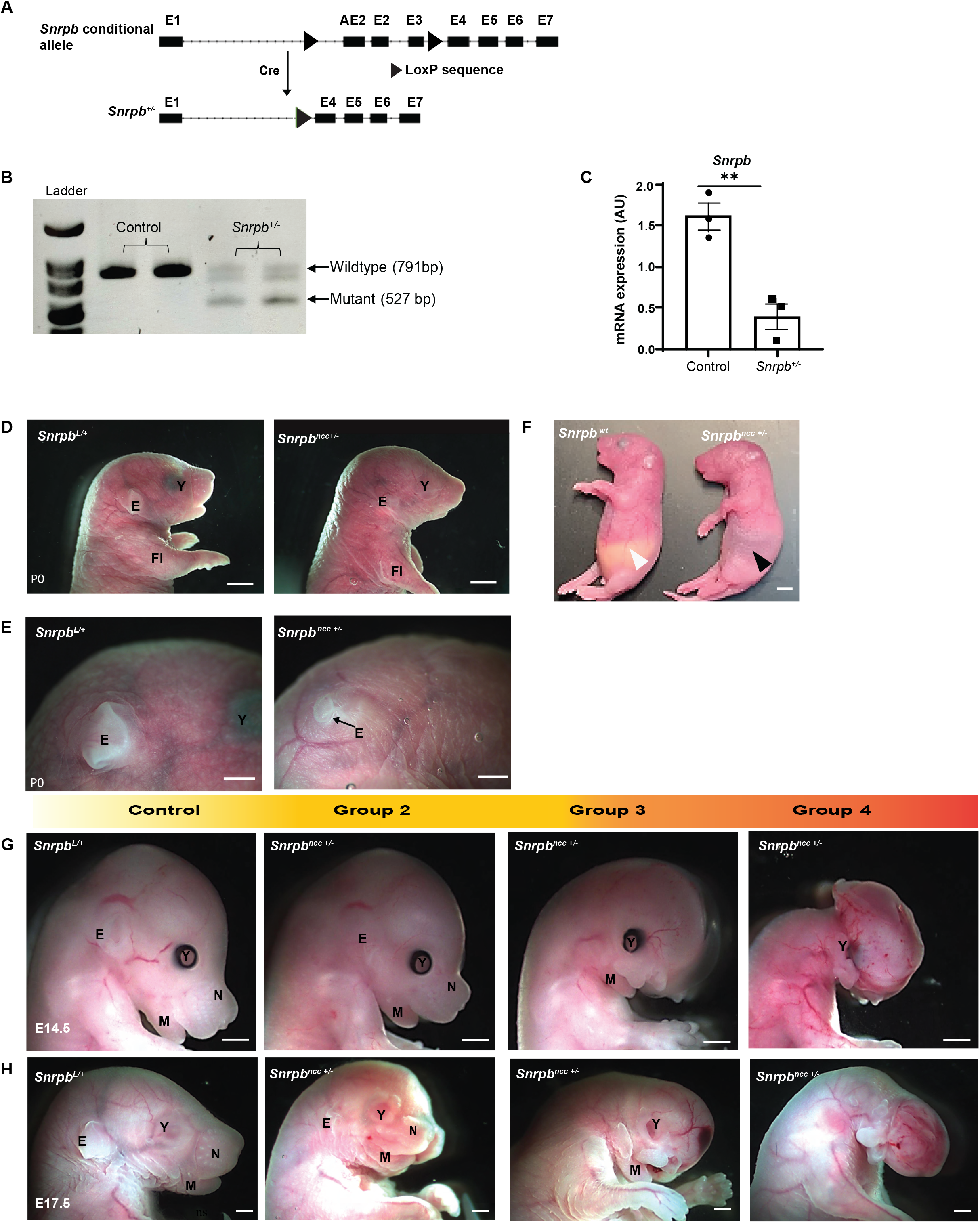
Deletion of Exons 2-3 of *Snrpb* results in reduced *Snrpb* level and craniofacial malformations of varying severity. **A**. Schema of the conditional *Snrpb^LoxP/+^* allele generated using CRISPR/Cas9 and the deletion generated in the presence of *Cre*. **B**. Deletion of the *LoxP* flanked exons 2 – 3 in *Snrpb* produces a shorter transcript of 527 base-pairs. **C**. RT-qPCR showing a significant decrease in *Snrpb* level in E8.5 heterozygous mutant embryos. **D**. P0 *Snrpb^ncc+/-^ (Snrpb^Loxp/+^*; *Wnt-1Cre2^tg/+^*) pups have an abnormally shaped head, micrognathia and abnormal outer ears. **E**. Higher magnification showing a hypoplastic pinna (E) in the *Snrpb^ncc+/-^* mutant (black arrow). **F**. *Snrpb^ncc+/-^* mutant pups lack milk spot in their stomach (black arrowhead), which is visible in the *Snrpb* wildtype littermate (white arrowhead). **G-H**. Representative images of *Snrpb* wild type (*Snprb^L/+^*) and *Snrpb^ncc+/-^* mutant embryos at E14.5 and E17.5. *Snrpb^ncc+/-^* mutant embryos show a range of craniofacial malformations, Group 2 had abnormal outer ear, cranial and mandibular hypoplasia, Group 3 had nasal clefts, Group 4 showed severe abnormalities including absence of the head and face. E=Ear, Fl=Forelimb, M=Mandible, N=Nose, Y=Eye,

### *Snrpb* is required in the neural crest cells for craniofacial development and postnatal survival

We next used a *Wnt1-Cre2* transgene to delete *Snrpb* in the neural tube and neural crest cells (*Snrpb^ncc+/-^*) to examine its role during craniofacial development. No *Snrpb^ncc+/-^* mutant pups were found at postnatal day (P)1 and P21 (p>.0001, chi-square). At P0, we recovered 5 heterozygous *Snrpb^ncc+/-^* pups from 6 litters (n=47). Most had no visible milk spots, indicating that they failed to feed (n=4/5). One *Snrpb^ncc+/-^* pup was morphologically normal, while the rest had abnormally shaped heads, short snouts, and small outer ears (n=4) (Fig. 1D-F). To determine when *Snrpb* is first required in neural crest cells for embryonic survival and craniofacial development, we collected and analyzed embryos from E9.0 and E17.5. *Snrpb^ncc+/-^* embryos are found at the expected Mendelian ratio until E17.5, when significantly fewer mutant embryos were found (Table 1A) (P<0.025, chi-square test). At E14.5 and E17.5, 43% and 25% of *Snrpb^ncc+/-^* embryos were dead and undergoing resorption, respectively. Thus, a significant number of *Snrpb^ncc+/-^* embryos die between E14.5 and birth.

**Table 1A.** Genotype of embryos collected from E9.0 – E17.5 from mating between *Snrpb^L/+^* and *Wnt1-Cre*^Tg/+^ mice. Dead embryos undergoing resorption are in parenthesis. *Chi square test indicated a significant difference from Mendelian expectation at E17.5 (P < .05).

| *Stage* | *Snrpb*<sup>L/+</sup><br>(resorbed) | *Wnt-Cre*<sup>Tg/+</sup><br>(resorbed) | *Snrpb*<sup>+/+</sup><br>(resorbed) | *Snrpb*<sup>L/+</sup>; *Wnt-Cre*<sup>Tg/+</sup><br>(resorbed) | *Total genotyped* |
| --- | --- | --- | --- | --- | --- |
| E9.0 | 16 | 16 | 18(1) | 23 (2) | 73 |
| E9.5 | 46 | 52(3) | 56 | 52(1) | 206 |
| E10.5 | 31(2) | 39 | 42 | 49(1) | 161 |
| E11.5 | 10 | 12 | 13 | 12 | 47 |
| E12.5 | 14 | 8 | 12 | 16 (2) | 50 |
| E14.5 | 17 | 19 | 16 | 21(9) | 73 |
| E17.5 | 37 | 20 | 27(1) | 16(4) * | 100 |
| **Total** | **171** | **166** | **184** | **189** | **710** |

We found that at E9.0, *Snrpb^ncc+/-^* embryos with thirteen or less somites, were indistinguishable from control (*Snrpb^+/+^* or *Wnt^tg/+^*) littermates (Table 1B; Fig. S1A). However, morphological abnormalities were first apparent in live *Snrpb^ncc+/-^* embryos at E9.5. At this stage 35% of *Snrp^ncc+/-^* mutants (n= 18), exhibited hypoplasia of the midbrain and hindbrain. At E10.5, 74% of mutant E10.5 (n=43) also showed hypoplasia of the frontonasal, maxillary, and mandibular prominences, the pharyngeal arches in addition to a smaller midbrain and hindbrain (Table 1B; Fig. S1 B-C). E11.5, *Snrpb^ncc+/-^* embryos (n= 12) could be sorted into three groups based on their shared phenotypes. We assigned the 17% of embryos, which were morphologically normal and indistinguishable from controls to Group 1/Normal (n=2); the 17% of mutants with hypoplasia of the developing brain, face and head were assigned to group 2 (n=2); and the remaining 66% were assigned to group 3 (n=8). Abnormalities found in group 3 included hypoplasia of the midbrain, swelling in the forebrain, sub-epidermal swelling, an absence of the frontonasal and the maxillary prominences, and a hypoplastic mandibular arch (Table 1B; Fig. S1D). At E12.5, 25% were morphologically normal (n= 4; group 1). Morphologically abnormal mutants at this stage were classified as group 2 (19%; n=3) or group 3 (25%; n=4), respectively. Mutants in group 2, exhibited clefts in the frontonasal prominence and the mandible, while those in group 3, had hypoplasia of the midbrain, an abnormal forebrain, and cleft of the hypoplastic frontonasal and maxillary prominences (Table 1B; Fig. S1E). A fourth phenotypic group comprising of 19% of *Snrpb^ncc+/-^* embryos was found (n=3) at E12.5. Embryos in this group showed absence of the ventral portion of the head and face, edema in the head, and a hypoplastic mandibular arch (Table 1B; Fig. S1E right panel). At E14.5, morphologically normal, group1 *Snrpb^ncc+/-^* embryos comprised 8% of live mutant embryos (n=1). Mutant embryos in group 2 (n=3), had a hypoplastic pinna, a dome-shaped head, and nasal clefts; and those in group 3 (n=4), showed hypoplasia and cleft of the frontonasal, maxilla and mandibular regions, and sub-epidermal edema (Table 1B; Fig. 1G, S1F). *Snrpb^ncc+/-^* embryos in group 4 (n=4) showed the most severe abnormalities (Table 1B; Fig. 1G, S1F), and were missing the ventral portion of the head and face. At E17.5 phenotypically normal group 1 embryos were not found. Half of the mutant embryos found alive were classified as group 2, (n=6) and the remainders were in groups 3 (n= 4) and 4 (n=2) (Table 1B; Fig. 1H, S1G). Thus, wild-type levels of *Snrpb* are critical in the neural crest cells from E9.5 onwards for normal development of the head and face.

**Table 1B.** The number of *Snrpb^ncc+/-^ (Snrpb ^L/+^;Wnt-Cre^Tg/+^*) embryos classified into each of four groups based on the presence and type of craniofacial malformations found. Embryos in group 1 where morphologically normal, whereas embryos in groups 2 to 4 had increasing phenotypic severity. * One litter was from mating between *Snrpb^L/+^* and *Wnt1-Cre* ^Tg/Tg^ mice.

| *Stage* | *Group 1* | *Group 2* | *Group 3* | *Group 4* | *Resorbed mutants* | *Total genotyped* |
| --- | --- | --- | --- | --- | --- | --- |
| E9.0 | 21 | 0 | 0 | 0 | 2 | 23 |
| E9.5 | 33 | 18 | 0 | 0 | 1 | 52 |
| E10.5* | 14 | 43 | 0 | 0 | 1 | 58 |
| E11.5 | 2 | 2 | 8 | 0 | 0 | 12 |
| E12.5 | 4 | 3 | 4 | 3 | 2 | 16 |
| E14.5 | 1 | 3 | 4 | 4 | 9 | 21 |
| E17.5 | 0 | 6 | 4 | 2 | 4 | 16 |

### *Snrpb* is required in neural crest cells for proper differentiation of both mesoderm and neural crest cells-derived cartilage and bones

Skeletal preparations with Alcian blue to stain cartilage and alizarin red to stain bone were used to examine development of neural crest cell derivatives in the developing head and face at E14.5, E17.5, and P0. Cartilage development in the head of the single E14.5 *Snrpb^ncc+/-^* group 1 embryo found was indistinguishable from that of controls (data not shown). However, in E14.5 and E17.5 *Snrpb^ncc+/-^* embryos belonging to groups 2 and 3 (n=8), the mesoderm-derived parietal bone and the intraparietal bone – which is derived from both mesoderm and neural crest cells – were hypoplastic (Fig. 2A, S2A-B). Neural-crest derived bones such as the temporal and alisphenoid bones were missing, while the frontal and nasal bones were hypoplastic (n= 5) (Fig. 2B, S2). Mutant embryos in these two groups also showed clefts of the nasal and pre-maxillary cartilages and bones, the palate, as well as hypoplasia of Meckel’s cartilage and its derivative, the mandible (Fig. 2C, 2D and S2). The zygomatic arch also failed to form in group 3 mutants (Fig. 2B). Those heterozygous mutants belonging to group 4 had hypoplasia of the basisphenoid bone and were missing neural crest cell-derived cartilages and bones that are normally found on the ventral surface of the head and face (Fig. 2B and S2). Furthermore, though the mandible formed in E14.5 and E17.5 *Snrpb^ncc+/-^* embryos in groups 2 and 3, it was both asymmetrical and bilaterally smaller when compared to that of controls (Fig. 2D, T-test, p<.0001). Distal ends of the jaws were abnormally shaped, while the proximal elements of the mandible such as the coronoid, condylar, and angular processes were not found in mutants (Fig. 2C). Additional defects found in mutant embryos included missing tympanic ring, hypoplasia or absence of the hyoid, and missing tracheal cartilages. In addition, ectopic cartilages and bones, which could not be conclusively identified (n=4 of 7) (Fig. 2E, S2C) were also found in the middle ear.

**Figure 2:**
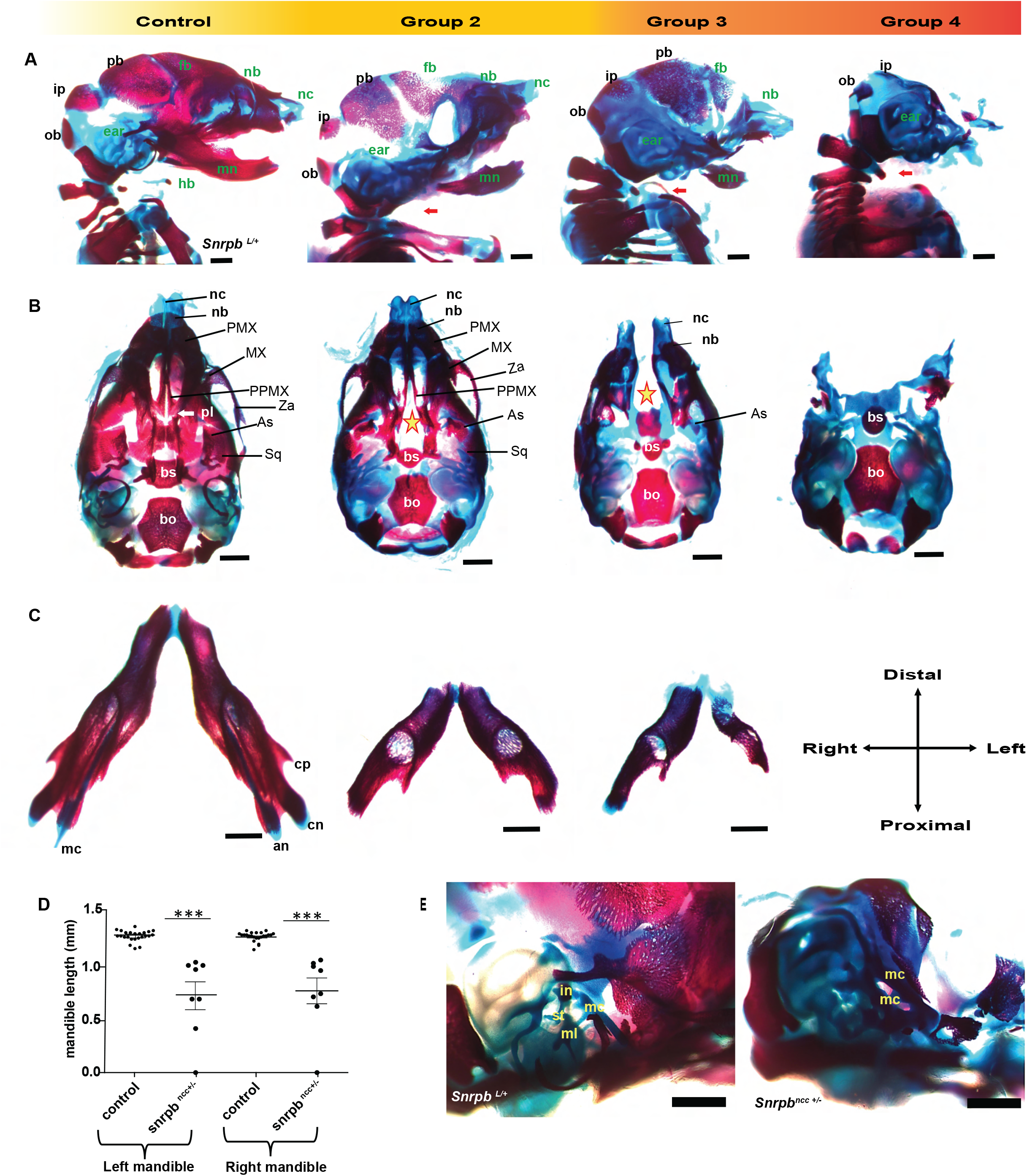
Craniofacial cartilages and bones of *Snrpb^ncc+/-^* embryos are hypoplastic or missing. **A and B**. Representative images of Alcian blue and Alizarin red stained E17.5 normal (*Snrpb^wt^* or *Snrpb^L/+^*) and *Snrpb^ncc+/-^* mutant embryos showing craniofacial abnormalities of varying penetrance. **A.** Sagittal view showing hypoplasia or absence of neural crest cell derived bones (labeled in green letters) in *Snrpb^ncc+/-^* mutants. The missing hyoid bone and tracheal cartilages are indicated with a red arrow. **B**. Ventral view of the skull showing palatal and maxillary clefts (star) in groups 2 and 3 *Snrpb^ncc+/-^* mutants, respectively, and the absence of the ventral craniofacial components in group 4 mutants. **C**. Representative images of the lower jaw of a normal and two *Snrpb^ncc+/-^* mutants showing asymmetric mandibles with no discernable angular, coronoid or condylar processes. **D**. Both left and right mandibles are significantly shorter in *Snrpb^ncc+/-^* embryos (p<.0001, T-test), when compared to controls (*Snrpb^wt^* and *Snrpb^L/+^* embryos). **E**. Representative images of higher magnification of the inner ear of a control and *Snrpb^ncc+/-^* embryo. Middle ear structures such as the stapes were absent in group 3 and 4 mutants, whereas a presumably duplicated Meckel’s cartilage and ectopic structures were found in a subset. an=angular process, As=alisphenoid bone, bo=basioccipital bone, bs=basisphenoid bone, cn=condylar process, cp=coronoid process, fb=frontal bone, hb=hyoid bone, in=incus, ip= intraparietal bone, mc=Meckel’s cartilage, ml=malleus, mn=mandible, MX=maxilla, nb=nasal bone, nc=nasal cartilage, ob=occipital bone, pb=parietal bone, PMX=premaxilla, pl=palatine, PPMX= palatal process of maxilla, Sq=squamous bone, st=stapes, Za=zygomatic arch

At P0, the morphologically normal *Snrpb^ncc+/-^* pup/group 1, had a curved but closed premaxilla (n=1; Fig. S3A). Furthermore, though 1 of the morphologically abnormal group 2 pup had a cleft in the premaxilla and was missing the palatine shelves (n=1), no bony palate defects were found in the remaining mutant pups (n=3) (Fig. S3A). Skull defects were found in both group 1 and group 2 *Snrpb^ncc+/-^* pups. These defects included reduced size of the squamous part of the temporal bone (n = 3), heterotopic ossification in the frontal suture (n=1), and a hypoplastic and asymmetric basisphenoid (n=3) (Fig. S3). Defects were also found in the mandible and middle ear. Meckel’s cartilage and the lower jaw which forms around it were asymmetric in most of these mutants (n=5). Specifically, the angular process was asymmetric in the group 1 mutant with 1 a wider angular process on one side (n=1) and three of the group 2 mutants (Fig. S3B). The articular surface cartilage was also absent or hypoplastic in group 1 and group 2 (n=2) mutants (Fig. S3B). Additionally, the condyloid and the angular processes were shortened bilaterally in one group 2 mutant (n=1). Middle ear defects such as absent, or abnormally shaped tympanic ring, and presence of ectopic ossification was found in group 1 and group 2 mutants (n=5; Fig. S3C). Our data indicate that *Snrpb* mutant neural crests can form cartilages but show deficiencies in ossification.

### Derivatives of cranial, cardiac, and trunk neural crest cells are abnormal in *Snrpb* mutants

In addition to cartilage and skeletal abnormalities, the dorsal root ganglia, and cranial nerve ganglion – which form from ectodermal placodes and neural crest cells – were also abnormal in *Snrpb^ncc+/-^* mutants. Neurofilament staining of group 2 E10.5 *Snrpb^ncc+/-^* mutant embryos showed that the cranial ganglia were reduced in size, and all had abnormal neuronal projection into the pharyngeal arches (n=2) (Fig. 3B). The ophthalmic branch of the trigeminal nerve (CN V) was reduced and did not extend over the lens, the maxillary projection appeared disorganized and missing, while the mandibular projection was reduced and appeared to have formed ventral to the first arch. In addition, an ectopic projection was found in CN V in mutants. Furthermore, the proximal portions of the geniculate (CN VII) and vestibulo-acoustic (CN VIII) ganglia were thicker than in *Snrpb^ncc+/+^*. Similarly, the glossopharyngeal nerve (CN IX) was abnormally thicker in the proximal region before the pharyngeal arch, had ectopic projection into pharyngeal arch 2, and reduced projection into pharyngeal arch 3. Finally, the proximal portion of the vagus nerve (CN X) was relatively normal but had an abnormal bundle at the distal end with reduced projections into the heart. Furthermore, the dorsal root ganglia which are derived from trunk neural crest cells were reduced in size and bifurcated at the proximal end (Fig. 3D).

**Figure 3:**
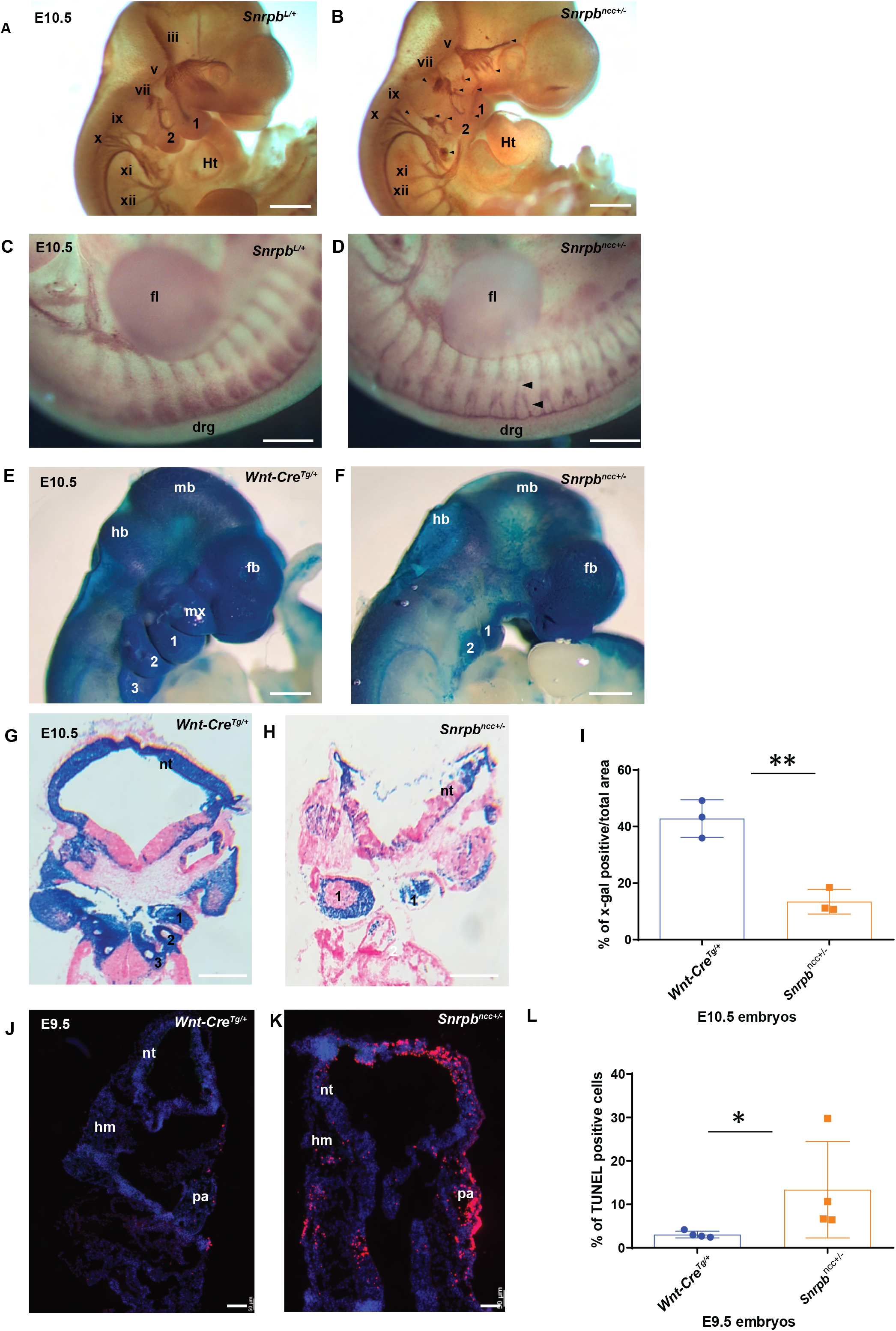
Abnormal cranial ganglia, reduced cranial neural crest cells, and a significant increase in cell death in *Snrpb^ncc+/-^* mutants. **A**., **B**. Representative images of E10.5 control (*Snrpb^L/+^*) and *Snrpb^ncc+/-^* group 2 embryos stained with 2H3. **B**. *Snrpb^ncc+/-^* mutants (n=2) showed abnormal projections of nerves to the pharyngeal arches (cranial ganglion v and vii) and heart (cranial ganglion ix) and absence and abnormal bundling of cranial nerves (all are shown in black arrowheads). **C.** When compared to controls (*Snrpb^L/+^*), the dorsal root ganglia are bifurcated and reduced in mutants (black arrowheads) shown in **D**. **E-H** Representative images of wholemount and sections of X-gal stained E10.5 control (*Wnt-Cre^tg/+^*) and *Snrpb^ncc+/-^* group 2 mutant embryos. *Snrpb^ncc+/-^* mutants show reduced X-gal staining in the craniofacial region and in the pharyngeal arches (n=4) when compared to the control embryos (n=4) **I**. Quantification of the area stained with X-gal showed a significant reduction in mutants (n=3) when compared to control littermates (n=3) (T-test, p<.005 **), Error bars indicate standard error of mean (SEM) **J.,K.**. Representative image of sections of TUNEL stained E9.5 control (*Snrpb^wt^* or *Snrpb^L/+^*) and *Snrpb^ncc+/-^* mutant embryos. **L**. Quantification showed an increase in the percent of TUNEL positive nuclei (red) in the craniofacial region of mutants (n=3) (T-test, p<.05 *), Error bars indicate standard error of mean (SEM). 1, 2, 3 =pharyngeal arches 1, 2 and 3, drg=dorsal root ganglia, fb=forebrain, fl=forelimb, hb=hindbrain, hm=head mesenchyme, ht=heart, mb=midbrain, mx=maxillary prominence, nt=neural tube, pa = pharyngeal arch

Micro CT scans of an E14.5, *Snrpb^ncc+/-^* embryo from group 4, revealed that the nasal septum and nasopharyngeal cavity were not formed, however, the oropharynx, tongue and pituitary gland were present (Fig. S4A). CT scans of an E17.5 wild type (control) and mutant embryos revealed that the aorticopulmonary septum, which is derived from cardiac neural crest cells, did not differentiate in the E17.5 group 2 *Snrpb^ncc+/-^* embryo (Fig. S4B-S4E). Furthermore, the thymus gland, a derivative of the third pharyngeal pouch, also failed to form in this mutant embryo (Fig. S4F). Finally, at all stages E12.5 (n=1), E14.5 (n=1) and E17.5 (n=1), the cerebral cortex was abnormally thin, and the lateral ventricles were enlarged in mutants (Fig. S4, and data not shown). Altogether, our morphological analysis indicates that *Snrpb* is required for the formation of structures that are derived from or induced by neural crests along the anterior-posterior axis. Our data also suggest that cardiac anomaly might contribute to death of *Snrpb^ncc+/-^* embryos and pups.

### Neural crest cells require wild-type levels of *Snrpb* for their survival in the craniofacial region

To track *Snrpb^ncc+/-^* heterozygous neural crest cells and their derivatives, we introduced the ROSA lacZ (Soriano P, 1999) and ROSA mT/mG (Mazumdar *et al*., 2007) reporters into our mutant line. At E9.5, wild-type and *Snrpb^ncc+/-^* embryos (morphologically normal (n=1) and abnormal (n=2)) showed a comparable proportion of X-Gal positive cells in the head and pharyngeal arches (Fig. S5A-C) with no statistically significant differences. Similarly, when the mT/mG reporter was used to visualize GFP positive Cre-expressing cells in control (n=3) and morphologically normal *Snrpb^ncc+/-^* mutants (n=3), no difference was found (Fig. S5D – H). However, at E10.5, morphologically abnormal group 2 mutant embryos (n=4) showed a reduced proportion of X-Gal positive cells in the head region (Fig. 3G-I), and this difference was statistically significant when compared to wild type (T-test, p=.003). To determine if reduced proliferation and/or increased apoptosis contribute to loss of X-Gal positive *Snrpb* heterozygous cells, E9.5 and E10.5 embryos were analyzed after Phosphohistone H3 immunostaining and TUNEL. No significant difference in proliferation was found between E9.5 control and *Snrpb^ncc+/-^* embryos (n=4; 3-group 1/normal and 1 group 2) and E10.5 (n=5; 1 group1/normal and 4 group 2) (Fig. S5 J-K). However, a statistically significant increase in TUNEL positive cells was found in the developing head region of E9.5 *Snrpb^ncc+/-^* embryos (T-test, p=.029) (n=4; 3-group 1/normal and 1 group 2) when compared to controls (Fig. 3J-L). Our data indicate that *Snrpb* heterozygous cells migrate into the developing head region and the pharyngeal arches. However, a subset of these cells undergoes apoptosis and are lost in mutant embryos by E10.5.

### Mutations in *Snrpb* cause an overall increase in both skipped exon and intron retention

To identify the molecular events that precede cell death, heads of E9.0 *Snrpb^ncc+/-^* embryos with 11-13 somite pairs, prior to morphological defects, were isolated and used for RNA sequencing analysis. Surprisingly, gene expression data did not reveal a major distinction between mutant and wild type embryos and the samples did not cluster by genotype. This was further corroborated by differential gene expression (DEG) analysis, which identified very few (76) DEGs: 50 upregulated and 26 downregulated in the mutant embryos. This low number of DEGs is consistent with the lack of a clear phenotypic distinction at this developmental stage. However, the DEGs which were identified could already be characterized into relevant molecular pathways, specifically belonging to the P53 signaling pathway and representing components of the spliceosome (Fig. 4A).

**Figure 4:**
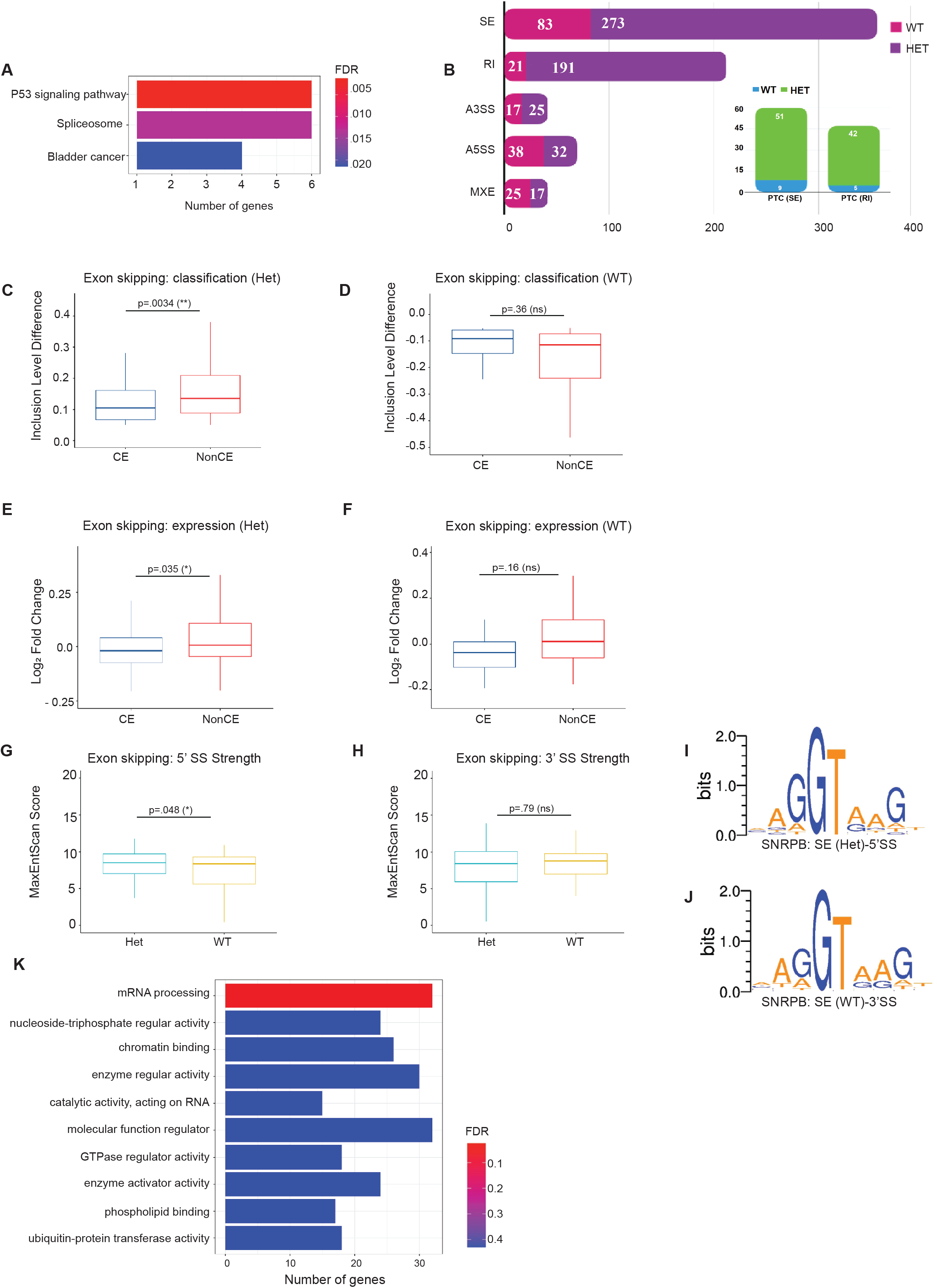
*Snrpb^ncc+/-^* mutant heads show aberrant splicing including both increased exon skipping and intron retention. **A**. DEGs identified in *Snrpb^ncc+/-^* (Het) mutants could be grouped into molecular pathways belonging to the p53 signaling pathway and the spliceosome **B**. A much larger number of transcripts was found to be abnormally spliced in *Snrpb^ncc+/-^* mutants when compared to controls (*Snrpb^wt^*). The most abundant differentially spliced events (DSE) were skipped exons (SE) and retained introns (RI). A strong tendency towards increased exon skipping and intron inclusion in the mutant samples was observed; there were more SE (273 in Het versus 83 in WT) and RI (191 in Het versus 21 in WT). DSE were predicted to introduce premature termination codons (PTC) more often in Hets when compared to wild type **C-F**. In Hets, Exon skipping was significantly higher for non-constitutive exons (NonCE) when constitutive (CE) vs (NonCE) was examined,T-test **G-J**. An analysis of Splice site (SS) strengths in DSE revealed significantly stronger 5’SS in mutants (P=.05, T-Test). **K**. Pathway analysis of genes with DSEs showed that they were strongly associated with mRNA processing.

In contrast to the low number of DEGs, a large number of transcripts were found to be abnormally spliced. We identified 722 significant (FDR = 0.1) differentially spliced events (DSE) between the *Snrpb^ncc+/-^* (Het) and wild-type (WT) samples. The most abundant of these DSE were skipped exons (SE) and retained introns (RI) (Fig. 4B). While the high proportion of SE could be expected based on prior alternative splicing studies, it was notable that more than 30% of the total AS events detected were RI events. We observed a strong tendency towards increased exon skipping and intron inclusion in the mutant samples; there were more SE (273 in Het versus 83 in WT) and RI (191 in Het versus 21 in WT) (Fig. 4B). Consistent with the absence of significant gene expression changes, DSEs in *Snrpb^ncc+/-^* embryos did not lead to significant changes in inclusion of PTC containing exons or introns (Fig. S6 A-D). However, though SEs were more likely to be alternative exons (non-constitutive) in heterozygous (p=0.0034) versus wild type (Fig. 4C-D), expression of transcripts with SE of constitutive exons was significantly reduced in mutants (p=0.0035) when compared to wild type (Fig. 4E-F). Those global trends in splicing are consistent with those previously found in cell culture, suggesting that SNRPB deficiency results in increased skipping of alternatively spliced exons resulting from reduced recognition of splicing signals (Correa *et al*., 2016).

### Increased skipped exon and intron retention in *Snrpb* mutants are not linked to identifiable sequence features

We next investigated whether the aberrant splicing events in the mutant embryos could in fact be characterized by specific sequence features. We compared alternative events preferentially found in the mutants to two control groups: 1) events preferentially found in the wild-type embryos, and 2) a set of 1000 randomly chosen alternative events. Specifically, we aimed to test whether aberrant events in the mutants were associated with weaker splice signals. While there was a very slight trend towards weaker splice site scores (MaxEntScan, Yeo *et al*., 2004) of RI in mutant as compared to *Snrpb* wild type embryos, the differences were small and not statistically significant (Fig. S6E-F). In contrast, 5’ SS strength was significantly higher in *Snrpb^ncc+/-^* heterozygous when compared to *Snrpb* wild type embryos (Fig. 4G), while the 3’ SS was comparable between mutants and *Snrpb* wild types (Fig. 4H). We also analyzed the strength and position of predicted branch point (BP) signals (LaBranchoR, Paggi and Bejerano, 2018), but again we did not find notable differences (Fig. S6K-N), with the exception of a slight preference for a more distal branch point location of mutant-specific SE events (27 bp in mutant versus 25 bp in the random set, p= 0.026). We also looked at general base composition in SE and RIs and no statistically significant difference was observed (Fig. S6O-R), though the GC content in retained introns was slightly increased (Fig. S6S-T). We also found no statistical differences in the length of SEs and RIs, although RIs were generally shorter in mutants when compared to wildtype (data not shown). Finally, we scanned for the frequency of RNA-binding protein motifs around the mutant specific events (rMAPS2, Jae Y Hwang *et al*. 2020), but did not identify significant enrichment of recognition signals of known splicing factors.

Overall, we did not find a compelling indication that the splicing aberrations present in mutants are linked to identifiable sequence features. The slight preference for stronger 5’ SS, branch point site location (BPS) and intronic nucleotide composition are notable but will need further scrutiny using more sensitive experimental designs. However, pathway analysis indicated that DSEs genes were significantly associated with mRNA processing (Fig. 4K). Thus, the relatively large number of splicing aberrations, as compared to differentially expressed genes, detected at this developmental stage supports the hypothesis that these general splicing defects precede aberrations in gene expression and initiate the molecular cascade that leads to phenotypic changes.

### Increased skipping of *Mdm2* exon 3 and *Mdm4* exon 7, key regulators of P53, are associated with increase nuclear P53 in heads of *Snrpb^ncc+/-^* embryos

We next investigated the key splicing changes that could explain craniofacial malformations in *Snrpb^ncc+/-^* embryos. We found increased skipping of exon 3 of *Mdm2* and exon 7 of *Mdm4*, regulators of the P53 pathway in our RNA-Seq analysis and confirmed these increases by RT-PCR (Fig. 5A-B). Increased skipping of these exons was previously reported in cultured *Snrpb* knockdown cells and shown to increase levels of nuclear P53 in mouse embryos with mutations in *Eftud2* a core component of the spliceosome (Beauchamp *et al*., 2021, Alstyne *et al*., 2018, Correa *et al*., 2016). Immunohistochemistry with an antibody to P53 revealed a significant enrichment of nuclear P53 in E9.5 mutant heads (Fig. S7A-B). Furthermore, levels of the P53-regulated genes *Trp53inp1, Ccng1*, and *Phlda3* were increased, and this increase was statistically significant when levels of *Ccng1*, and *Phlda3* were compared between E9.0 *Snrpb* wild type and mutant embryos, but not at E9.5 (Fig. 5C-D). Thus, we conclude that the increased exon skipping in *Mdm2* and *Mdm4* results in increased nuclear P53, and levels of P53 target genes in *Snrpb^ncc+/-^* embryos, prior to morphological abnormalities. Since P53 activation can lead to increased apoptosis, we postulate that increased P53 activity contributes to apoptosis of *Snrpb^ncc+/-^* mutant cells.

**Figure 5:**
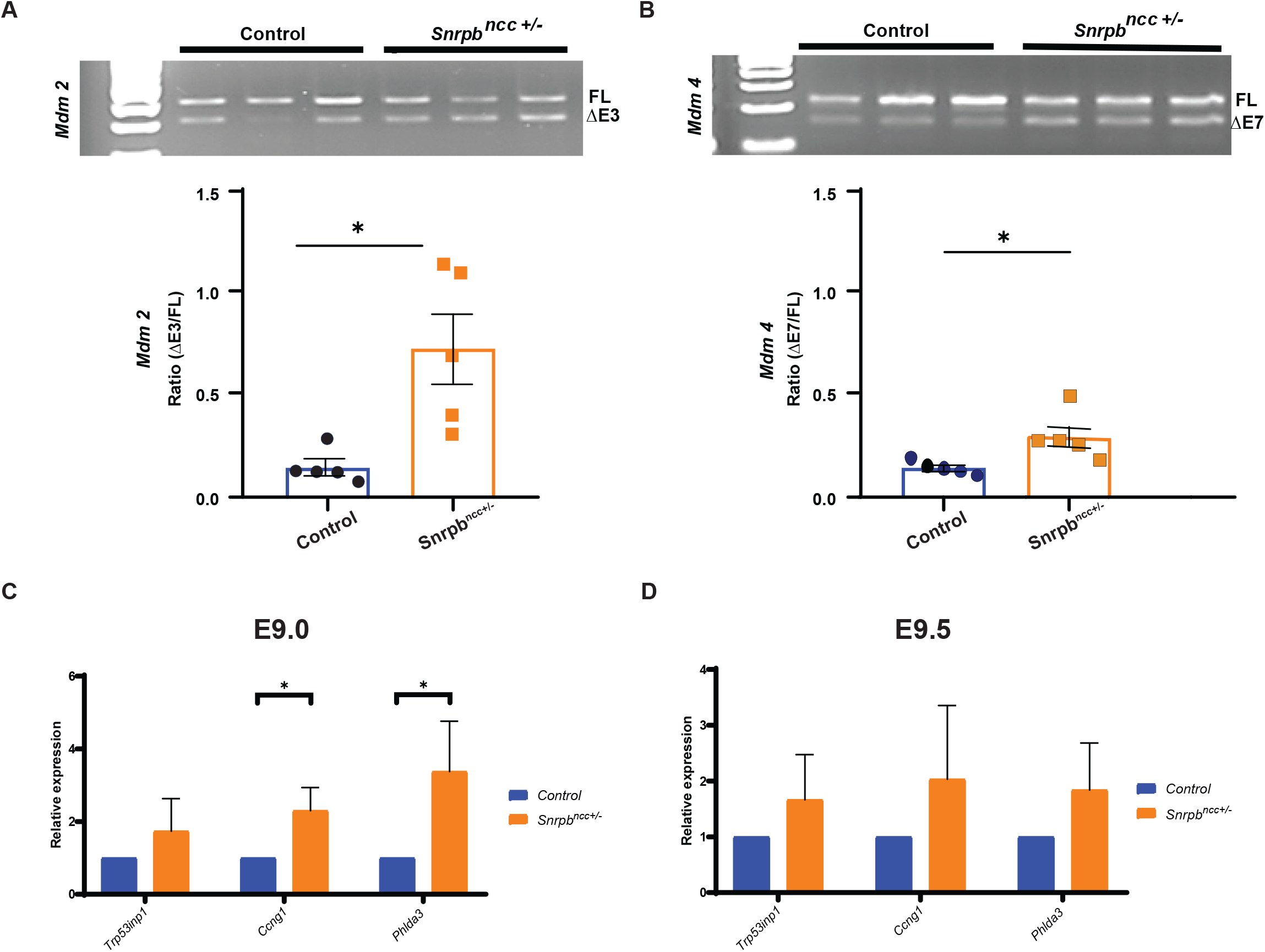
Increased exon skipping in two regulators of P53; *Mdm2* and *Mdm4*, and significant increases in P53-target genes in *Snrpb^ncc+/-^* mutant heads. **A**. and **B**. Representative images of RT-PCR showing long (FL) and short transcripts (ΔE) produced in *Mdm2* and *Mdm4* in E9.0 control (*Snrpb^wt^* or *Snrpb^L/+^*) and *Snrpb^ncc+/-^* embryos. Quantification revealed a significant increase in the ratio of *Mdm2* and *Mdm4* transcripts containing a skipped exon over the longer transcript in mutants (t-test, p<.05), Error bars indicate standard deviation (SD). **C**. Levels of P53 target genes were significantly increased in morphologically normal E9.0 *Snprb^ncc +/-^* embryos (n=5) when compared to controls (n=5) (t-test, p<.05), **D.** At E9.5 though levels of P53 target genes were increased in *Snrpb^ncc+/-^* embryos (n= 3 group1 and n=2 group 2), this difference was not significant, Error bars indicate standard error of the mean (SEM). FL= full length transcript, ΔE3= transcript with exon 3 skipped, ΔE7= transcript with exon 7 skipped.

### Reducing levels of P53 does not rescue craniofacial defects in *Snrpb^ncc/+^* embryos

We next tested if reducing levels of P53 prevents craniofacial malformations in *Snrpb^ncc+/-^* embryos. We crossed *Trp53^loxP/+^; Wnt1-cre2^tg^* mice and *Snrpb^loxP/+^* mice and collected E10.5 and E17.5 *Snrpb^ncc+/-^; P53^ncc+/-^* double heterozygous embryos for analysis. We found no significant difference in the proportion of *Snrpb^ncc+/-^; P53^ncc+/-^* embryos with mild to severe craniofacial defects (n=4), when compared to *Snrpb^ncc+/-^* mutants (n=3/3) (Fig. S7C-D and not shown). We then generated *Snrpb^ncc+/-^* with two mutant *Trp53* alleles in their neural crest cells (*Snrpb^ncc+/-^; P53^ncc-/-^*) for cartilage and skeletal analysis. E14.5 *Snrpb^ncc+/-^; Trp53^ncc-/-^* mutant embryos (n=2) resembled *Snrpb^ncc+/-^* mutants found in group 2 (Fig. S7E). Similarly, E18.5 *Snrpb^ncc+/-^; Trp53^ncc-/-^* mutant embryos (n=4) were morphologically similar to group 2 *Snrpb^ncc+/-^* mutants; they had microcephaly, a shorter snout and micrognathia (Fig. S7G). Cartilage and skeletal preps revealed reduced ossification of the frontal bone, cleft palate, and asymmetric and abnormal development of the lower jaw (Fig. S7F, H-J). Although we did not find any *Snrpb^ncc+/-^* embryos with homozygous loss of *Trp53* in their neural crest cells that were severely abnormal, we did not recover any *Snrpb^ncc+/-^; Trp53^ncc-/-^* pups when the matings were allowed to go to term or find any morphologically normal mutants, suggesting that removing both copies of *Trp53* does not rescue craniofacial defects in *Snrpb^ncc+/-^* embryos.

### Genes required for craniofacial development are abnormally spliced in *Snrpb ^ncc+/-^* mutants

To identify additional abnormal splicing events which could explain craniofacial malformations in *Snrpb ^ncc+/-^* embryos, we queried the MGI database to determine if any transcripts with statistically significant DSEs were required for craniofacial development (Bogue *et al*., 2017). We identified 13 transcripts required for craniofacial development or stem cell development with significant increases in exon skipping (Table S1). Increased exon skipping in 5 of these genes: *Pdpk1, Rere (Atr2), Mcph1, Nf1*, and *Dyrk2*, is predicted to introduce a pretermination codon. The remaining exon skipping events are not predicted to result in PTC but may alter gene expression and/or function. In fact, all except one of these DSEs were in constitutive exons. We then queried our RNAseq dataset to determine if the expression level of these genes was altered in *Snrpb* mutants. We found no significant changes in levels of transcripts with PTC or Non-PTC skipped exons. We then selected 3-transcripts and performed RT-PCR to confirm that the skipped exon events identified in the RNAseq analysis were present in *Snrpb* mutant heads (Fig. 6). This analysis revealed presence of transcripts with skipped exons for *Smad2, Pou2f1* and *Rere*, although the percent spliced events for *Smad2* and *Pou2f1* were below 10% and not significant when ratio of short/long transcript in control and mutant was compared. We postulated that abnormal increases in exon skipping in these 13-transcripts, which are required for normal craniofacial development, may contribute to craniofacial defects in *Snrpb^ncc/+^* mutants.

**Figure 6:**
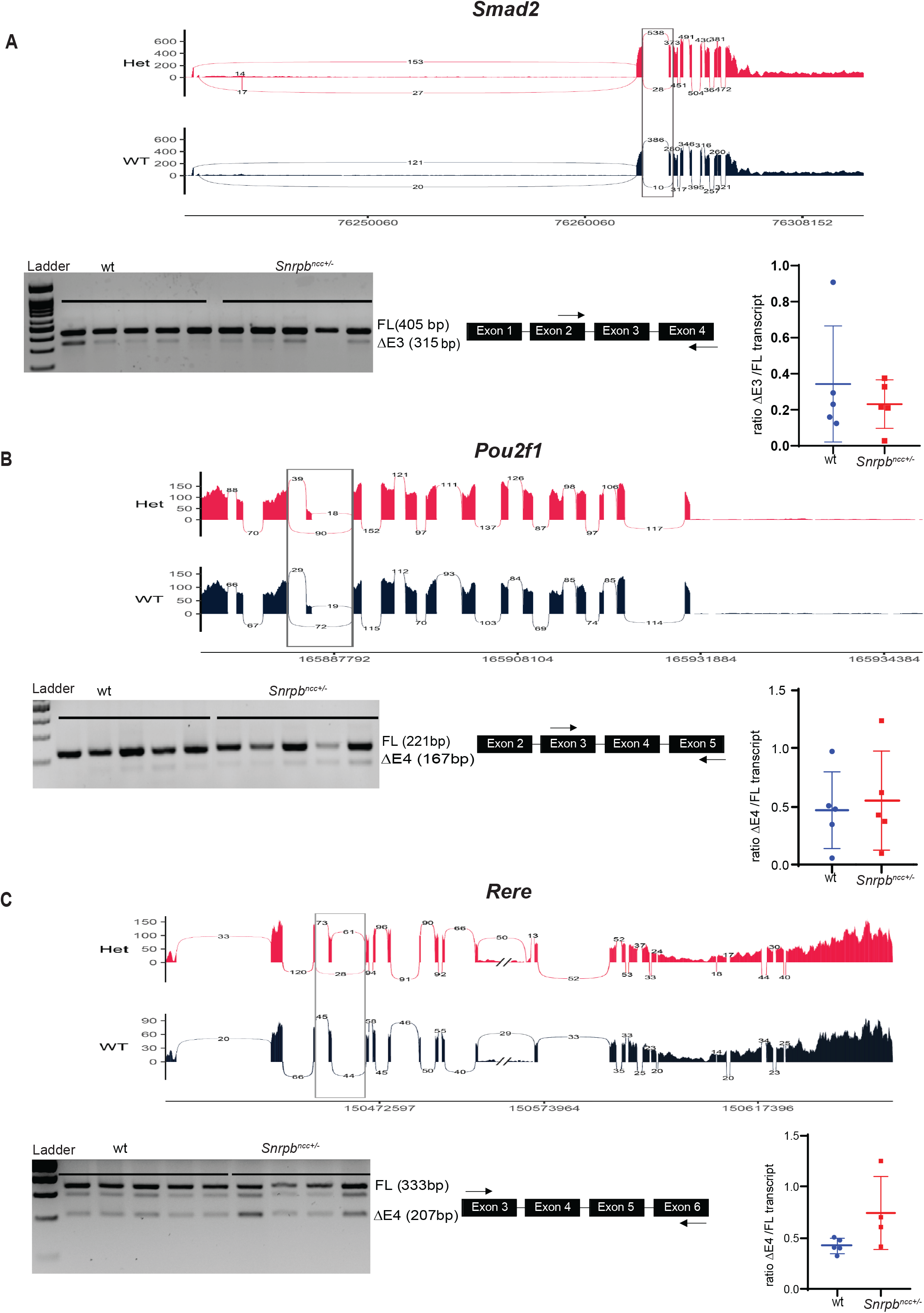
Transcripts with exon skipping events are found in heads of E9.0 control and mutant embryos. Sashimi plot for the exon skipping events found for **A**. *Smad2*, **B***. Pou2f1* and **C**. *Rere*. Under each sashimi plot, representative gel for RT-PCR showing the presence of transcripts with the predicted exon skipping event. The location of primers used to amplify transcripts is shown on the right. No significant difference was found in the ratio of short/long transcripts (T-test). Error bars in the graphs indicate standard error of the mean (SEM). FL=Full length, ΔE=skipped exon.

### Wild-type levels of *Snrpb* are required for normal expression of *Fgf8, Shh* and *Msx2*

The surface cephalic ectoderm, of which the facial ectoderm zone (FEZ) is a subregion, is essential for integrating proper growth of the craniofacial skeletons and brain and patterning the underlying neural crest (Griffin *et al*., 2013). The severe malformations found in the face and brain of *Snrpb* mutants suggest that the FEZ might not have formed. Therefore, we used *in situ* hybridization to examine expression of *Shh* and *Fgf8*, which are expressed in surface cephalic ectoderm and together help to define the FEZ (Griffin *et al*., 2013). At E9.5 before the FEZ forms, *Shh* was expressed in the ventral-most region of the neural tube, the floor plate, as well as the ventral prosencephalon of *Snrpb* wild type and *Snrpb^ncc+/-^* embryos (Fig. 7A, n=4; 3 group 1 and 1 group 2). At this stage, *Fgf8* was expressed in the mandibular epithelium, the frontonasal prominence, and the midbrain/hindbrain junction of control and *Snrpb^ncc+/-^* mutant embryos. However, the expression domain of *Fgf8* was abnormally expanded at these sites (n=4; 2 group 2 and 2 group 1) (Fig. 7B). Furthermore, expression of *Msx2*, a downstream target of *Fgf8* in the underlying neural crest cells (Griffin *et al*., 2013) was also abnormal in E9.5 *Snrpb^ncc+/-^* embryos. In *Snrpb^ncc+/+^* embryos (n=6), *Msx2* was expressed in the distal region of pharyngeal arches 1 and 2 (Fig. 7C). However, in *Snrpb^ncc+/-^* embryos, *Msx2* expression was abnormally extended proximally in these arches (n=4; 3 group1 and 1group 2) (Fig. 7C). These *in situ*s revealed that normal levels of *Snrpb* is required in the neural crest cells to restrict expression of *Fgf8* and its downstream target *Msx2* in the developing head and face in both morphologically normal and abnormal embryos.

**Figure 7:**
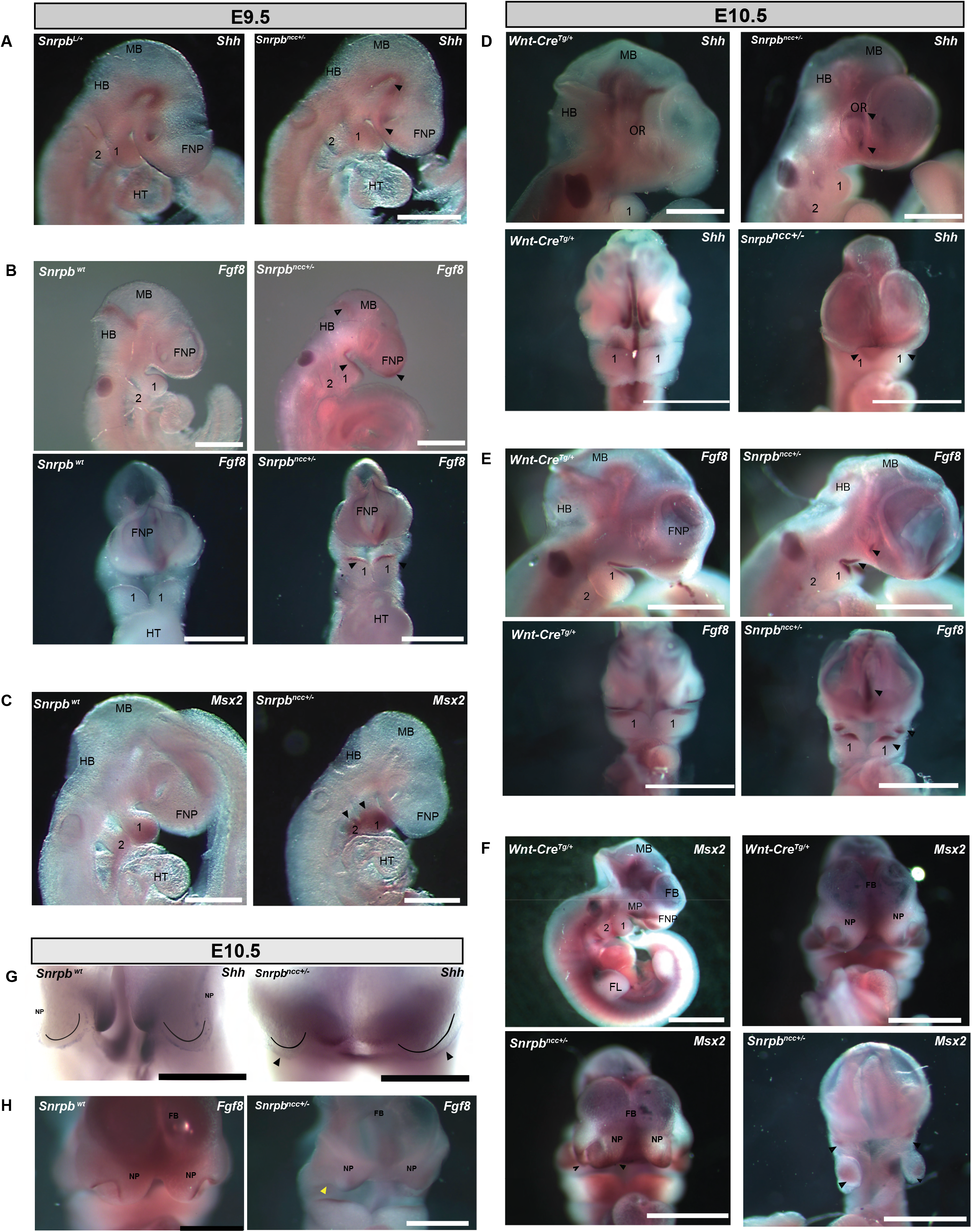
*Shh, Fgf8* and *Msx2* are mis-expressed in *Snrpb*^ncc+/-^ embryos. Representative images of E9.5 and E10.5 control (*Wnt-Cre2 ^Tg/+^* or *Snrpb^wt^*) and *Snrpb^ncc +/-^* embryos after wholemount *in situ* hybridization (ISH) to detect mRNA expression of *Shh, Fgf8* and *Msx2* **A**. *Shh* at E9.5 was expressed in the ventral-most region of the neural tube, the floor plate, as well as the ventral prosencephalon of control and *Snrpb^ncc+/-^* embryos (black arrowheads). **B**. Lateral (Top), and ventral (bottom) views of embryos showing that expression of *Fgf8* was expanded in the mandibular epithelium, the frontonasal prominence, and the midbrain/hindbrain junction in group 2 (black arrowhead) (n=4) mutant embryos. **C**. *Msx2* expression was extended proximally in pharyngeal arches 1 and 2 in group 1 mutant embryos (n=2; black arrowheads). At E10.5, wholemount ISH for **D**. Top panel, lateral view of embryos showing *Shh* expression in the diencephalon and the ventral forebrain in control and a group 2 mutant embryo. Ectopic expression was found in the dorsal and ventral optic lens (arrowheads) of the mutant. Bottom panel, ventral view showing expression of *Shh* on the oral ectoderm in control and group 2 mutant embryos (black arrowhead). **E**. Top panel shows representative images of lateral views of a control embryo, in which *Fgf8* was detected on the surface ectoderm of the mandible, maxillary and frontonasal prominences (n=3). In the representative group 2 *Snrpb^ncc+/-^* mutant embryo *Fgf8* expression on the mandibular ectoderm and in the region where the maxillary prominence would normally form (black arrowheads, n=3), ectopic expression was also detected in the lens. Bottom panel shows frontal view of reduced *Fgf8* expression in the hypoplastic lateral nasal prominence in mutants, and ectopic expression of *Fgf8* on the surface ectoderm of the medial nasal process, towards the midline (black arrowhead). **F**. Top panel shows a lateral and ventral view of *Msx2* expression in E10.5 control embryos. Bottom panel shows that in E10.5 group 1, *Snrpb^ncc+/-^* mutants on the left, *Msx2* was expressed in the lateral and medial nasal prominences, although expression appeared reduced and ventrally expanded in the medial frontal nasal region. On the right, in a group 2 *Snrpb^ncc+/-^* mutant with a hypoplastic frontonasal and maxillary prominences, *Msx2* was expressed in the region where the maxillary prominences would form and in the mandibular region of the hypoplastic first arch. **G**. and **H**. Higher magnification of the FEZ region of E10.5 embryos showing **G**. *Shh* expression in the developing mandibular periderm, and bilaterally on the surface ectoderm of the medial nasal prominences of control and group 2 *Snrpb^ncc+/-^* embryos (arrowheads). **H**. *Fgf8* expression in the lateral nasal prominence was reduced in group 2 mutants (yellow arrowhead), while ectopic expression of *Fgf8* was found on the surface ectoderm of the medial nasal process, towards the midline. 1, 2=pharyngeal arches1 and 2, FNP=frontonasal process, HB=hindbrain, HT=heart, MB=midbrain, MP=mandibular process, NP=nasal process, OR=optic region

In E10.5 group 2 *Snrpb^ncc+/-^* mutant embryos missing the frontonasal prominence, expression of *Shh* was found in the diencephalon and the ventral forebrain (n=3; Fig. 7D). Furthermore, ectopic *Shh* expression was found in the dorsal and ventral optic lens (arrows in Fig. 7D). In group 1 mutant embryos, the lateral and medial nasal processes were further apart than in wild type embryos but, *Shh* expression was found in the developing mandibular periderm, and bilaterally on the surface ectoderm of the medial nasal prominences (arrows in Fig 7G, n=1). In E10.5 wild type embryos, *Fgf8* was expressed on the surface ectoderm of the mandible, the maxillary prominences, and frontonasal prominences. In group 2 *Snrpb^ncc+/-^* mutant embryos with a hypoplastic frontonasal and maxillary prominences, expression of *Fgf8* was found on the mandibular ectoderm and in the region where the maxillary prominence would normally form (n=2) (Fig. 7E). In group 1 E10.5 *Snrpb^ncc+/-^* embryos, *Fgf8* expression in the lateral nasal prominence was reduced, while ectopic expression of *Fgf8* was found on the surface ectoderm of the medial nasal process, towards the midline (n=3, Fig. 7H). Similarly, in E10.5 group 2 *Snrpb^ncc+/-^* mutants missing the frontonasal and maxillary prominences, *Msx2* expression was expressed in the maxillary and in the mandibular region of the hypoplastic first arch (n=3). In group 1 *Snrpb^ncc+/-^* mutants, *Msx2* was expressed in the lateral and medial nasal prominences, although expression appeared reduced but ventrally expanded in the medial frontal nasal region (n=2; Fig. 7F). Thus, in *Snrpb^ncc+/-^* embryos where the lateral and medial nasal prominences formed, reduced expression of *Fgf8* in the ectoderm results in abnormal expression of *Msx2* in the underlying neural crest. We postulate that DSEs in genes important for midface development lead to abnormal expression of *Shh, Fgf8* and *Msx2* expression and mis-patterning of the developing craniofacial region.

## DISCUSSION

Splicing is an essential and ubiquitous process that generates mature mRNAs and increases the number and diversity of proteins from the genome (Chen and Manley, 2009; Nilsen and Graveley, 2010). SNRPB is an essential protein that facilitates assembly of the snRNPs proteins that carry out splicing. Surprisingly, mutations that increase levels of a non-functional *SNRPB* transcript result in CCMS, a craniofacial spliceosomopathy that is also associated with rib defects (Lynch *et al*., 2014; Barcot *et al*., 2015 and Beauchamp et al., 2020). To examine the role of *SNRPB* in the development of tissues affected in patients, we first designed guide RNAs and repair templates to generate a mutation in alternative exon 2 that would model those found in CCMS patients (data not shown). However, this strategy did not prove to be fruitful as we did not recover mutant mouse lines carrying this mutation (data not shown). Therefore, we generated a conditional mutant mouse line to test if wild-type levels of *Snrpb* were required for normal development. Herein, we showed that *Snrpb* is haploinsufficient in mice and required for normal splicing of key regulators of P53 and transcripts required for normal craniofacial development as well as expression of *Fgf8* and *Shh*. We show that morphological defects in *Snrpb* mutants were not associated with significant changes in gene expression but with disruptions in alternative splicing and patterning of the craniofacial region. We suggest that altered transcript ratios and expression of genes important for patterning the craniofacial region are responsible for malformations and embryonic death of mutant embryos.

In our conditional mutant mouse line, deletion of exon 2, alternative exon 2 and exon 3 in the presence of Cre is predicted to generate a shorter *Snrpb* transcript of 527 bp that encodes for a non-functional protein. When *β-Actin Cre* was used to delete the *loxP*-flanked exons, the resulting *Snrpb* heterozygous embryos died post-implantation. Therefore, we were unable to study the role of SNRPB in head and rib development in these mutants. Though further studies are needed to determine if there is a general growth defect or other roles for *Snrpb* at these early stages of development, our study indicates that heterozygosity for a loss of function allele of *Snrpb* is lethal. In fact, a single patient carrying a mutation in the 5’ UTR of *SNRPB* that was predicted to result in a null allele was more severely affected and failed to survive gestation (Lynch *et al*., 2014). Thus, our data support the hypothesis that *SNRPB* mutations commonly found in CCMS patients are not null mutations (Lynch *et al*., 2014).

To study the role of SNRPB in craniofacial development we used the *Wnt1-Cre2* transgenic mouse line to generate embryos with heterozygous mutation of *Snrpb* in their neural tube and neural crest cells. In *Snrpb^ncc+/-^* mutant embryos, craniofacial structures derived from neural crest cells, such as the nasal bone, palates, maxilla, mandible, and middle ear structures, are abnormally formed. Intriguingly, these structures are also commonly reported to be affected in CCMS patients, suggesting that abnormal neural crest cell survival and/or differentiation are responsible for defects in those patients. We also uncovered an absence or reduction of the hyoid bone and ectopic cartilages and bones that we could not identify in some *Snrpb^ncc+/-^* mutants. Similarly, accessory ossicles in the hyoid bone were found after CT scan of two CCMS patients by Tooley *et al*., 2016. We postulate that SNRPB is required in all neural crest cells, and their derivatives, a fact that is supported by the reduced or absent aorticopulmonary septum that is found in the microCT scan of *Snrpb^ncc+/-^* mutants. Furthermore, we propose that aorticopulmonary septal defects and or palatal clefts such as the ones found in E17.5 mutants contribute to death of *Snrpb^ncc+/-^* embryos, as was found in *Eftud2^ncc-/-^* mutants (Beauchamp *et al*., 2021). Finally, although phenotypes found in CCMS patients strongly suggested a requirement of SNRPB for endochondral ossification, our data show abnormal development of bones formed via both endochondral and intramembranous ossification, indicating an early role for SNRPB in skeletal development.

P53 stability and activity are known to be upregulated in response to mutation or disruption in the level of splicing factors (Alstyne *et al*., 2018, Correa *et al*. and 2016, Lei *et al*. 2020). In fact, we found increased skipping in two P53-regulators, *Mdm2* exon 3 and *Mdm4* exon 7, increased nuclear P53 and upregulation of P53 target genes in heads of E9.0 *Snrpb^ncc+/-^* mutants. In zebrafish and mouse, increased P53-activity contributes to craniofacial defects and knocking down or removing P53 genetically reduced apoptosis and improved development (Li L. *et al*., 2017; Mao H. *et al*., 2016). Additionally, we showed that the P53-inhibitor, Pifithrin-α, improved head and brain development in embryos with mutation of *Eftud2* in the neural tube and neural crest (Beauchamp *et al*., 2021). However, reducing or removing P53 genetically in the neural crest cells did not show rescue of craniofacial defects in *Snrpb^ncc+/-^* mutant embryos. Although the variable expressivity found in *Snrpb^ncc+/-^* embryos makes it difficult to rule out a partial rescue, our findings indicate that P53 alone is probably not responsible for the malformations that we found.

RNAseq is a sensitive method for examining gene expression (Wang, Z. *et al*., 2009), and our data indicate that reduced expression of SNRPB in mutant neural crest disrupts splicing and expression of genes important for craniofacial development. In fact, our RNAseq analysis using the head of morphologically normal E9.0 *Snrpb^ncc+/-^* embryos revealed many more DSEs than DEGs. We confirmed differential splicing of the P53 regulators, *Mdm2* and *Mdm4* which may lead to increased apoptosis and identified 13 transcripts important for craniofacial development that were abnormally spliced in *Snrpb^ncc+/-^* embryos. DSE may perturb gene expression levels, for example with the introduction of pretermination codon, or alter the activity or localization of the resulting gene product. For example, the DSE event associated with exon 3 of *Smad2* is predicted to increase the proportion of transcripts that encodes a much more potent effector of TGFβ/Nodal than the full-length SMAD2. This shorter protein heterodimerizes with SMAD3 to regulate many developmental processes, including growth of the mandible (Dunn *et al*., 2005). Similarly, increased skipping of exon 8 of *Ror2* may disrupt the ability of this receptor to interact with *Wnt5* during midface, ear, and jaw development (Schwabe, G.C. *et al*., 2004). Furthermore, deletions of constitutive exons may change the open reading frame, insert a pretermination codon, or generate an unstable transcript that is removed by nonsense-mediated decay. Hence, we predict that an increase in the proportion of transcripts with a missing constitutive exon will reduce levels of the associated proteins. Abnormal expression of *Fgf8* which is regulated by SMAD2 (Liu *et al*., 2004) and *Rere* (Kumar and Duester, 2014) may be one of the consequences of mis-splicing. Nonetheless, reduced migration of neural crest cells into the frontonasal region could also explain abnormal expression of *Fgf8* and *Shh*. Though we found no significant differences in the number of neural crest cells in heads of E9.0 control and *Snrpb* mutant embryos, reduced levels of *Nisch*, which binds to integrins to block cell migration (Ding *et al*., 2008), may disrupt migration of a specific subset of neural crest cells that cannot be identified with our current techniques. In the future, we will investigate contribution of DSEs to abnormal expression of *Fgf8* and *Shh* and to craniofacial defects in *Snrpb* mutants.

If, as we postulate, malformations in *Snrpb* mutants are due to DSE that leads to increased cell death and disruption of multiple pathways important for patterning, the variable penetrance found in mutants may reflect the proportion of cells that undergo cell death and the level of disruption in patterning. Thus, embryos where a large number of cells die would have absent craniofacial structure formations, and resemble group 3 or 4, whereas fewer cell death would lead to mutants classified as group 1/2. Furthermore, those in group 1 and 2, the severity of craniofacial malformation would then depend on the level of DSEs in those genes critical for patterning of the region. The absence of group 3 or 4 *Snrpb;Trp53* mutants support this hypothesis. Loss of P53 may rescue cell death and allow for development of craniofacial structures in these mutants. However, DSEs in patterning genes are presumably independent of P53 and may lead to malformations and embryonic death. Future characterization of cell death and patterning in *Snrpb;Trp53* double mutant embryos along with RNAseq experiments using morphologically normal and abnormal mutant heads may allow us to tease out these different contributors to craniofacial malformation and aid in identifying DSEs and pathways regulated by Snrpb.

Our working model is that dysregulation in the level of SNRPB, even if modest - as is likely the case with inclusion of the PTC containing alternative exon 2, perturbs the efficiency of splicing at the level of spliceosome assembly. Furthermore, although cells with reduced levels of *Snrpb* have an increased propensity to undergo apoptosis, increased DSE is found before they die. Therefore, we propose that splicing changes in important developmental genes, the proportion of cells that undergo apoptosis, and the timing of apoptosis may all contribute to the variable expressivity found in *Snrpb* heterozygous mice and in CCMS patients. In conclusion, we believe that our work using the first CCMS animal model shows evidence for both ubiquitous and development-specific roles of *Snrpb* during morphogenesis and provides much needed insights into the role of this splicing factor during embryogenesis.

## ACKNOWLEDGEMENT

We would like to thank Dr. Mitra Cowan, Platform Manager, McGill Integrated Core of Animal Modeling (MICAM) for performing the microinjection experiments. We would also like to thank members of the Majewska lab and Drs. Richard Behringer, and Jennifer Fish for their helpful comments on the manuscript. This project was funded in parts by Canadian Institutes of Health Research (CIHR), bridge funding from the Research Institute of McGill University Health Centre (RI-MUHC) and the Azrieli Foundation. We thank the Queen Elizabeth Scholars (QES) Program, the Research Institute of McGill University Health Centre (RI-MUHC) and the McGill Faculty of Medicine and Health Sciences for supporting Sabrina Alam. The funders had no role in study design, data collection and analyses, decision to publish or preparation of the manuscript. We acknowledge the professional and technical support from the Animal Resource Division (ARD) of RI-MUHC for maintaining our mice colonies. We are also thankful to the Small Animal Imaging Labs, Centre for Translational Biology, RI-MUHC for their support in performing the microCT scans. LJM and JM are members of the Research Centre of the McGill University Health Centre which is funded in parts by FRQS.

## AUTHOR CONTRIBUTIONS

L.J.M. and J.M. conceptualized and supervised the project; E.B., R.P. and S.J.Z. analyzed all the RNA sequencing data, S.K. performed RT-qPCR experiments and analyses; M.C.B performed 2H3 immunostaining, P53 immunohistochemistry, and in situ experiments and analysis, A.B. performed proliferation and apoptosis analysis; N.N. performed skeletal preparation analysis, S.S.A. performed all other experiments done in the study; L.J.M., J.M, and S.S.A wrote the manuscript with inputs from all authors.

## METHODS

### Resource Availability

#### Lead contact

Further information and requests for resources and reagents should be directed to and will be fulfilled by the Lead Contact, Loydie A. Jerome-Majewska.

#### Materials Availability

All antibodies, chemicals and most mouse lines used in this study are commercially available. All other unique materials are available upon request.

#### Data and code availability

Accession IDs provided by GEO: GSE180546, GSM5464627, GSM5464628, GSM5464629, GSM5464630, GSM5464631, GSM5464632

## METHOD DETAILS

### Mouse lines

All procedures and experiments were performed according to the guidelines of the Canadian Council on Animal Care and approved by the animal Care Committee of the Montreal Children’s Hospital. Wild-type CD1, mT/mG *Gt(ROSA)26Sor^tm4(ACTB-tdTomato,-EGFP)Luo^/J*, (Soriano, 1999), *Wnt1-Cre2* (Lewis *et.al, 2013*), and *β-Actin-cre* (Lewandowski and Martin, 1997) mice on the C57BL/6J genetic background were purchased from The Jackson Laboratory. The *R26R* strain (*Gt (ROSA)26Sor^tm1Sor^* (Mazumdar et al., 2007) on the mix C57BL/6J;129/S4 genetic background was a kind gift from Dr. Nagano. All strains were maintained on the CD1 genetic background. The *Trp53^tm1brn^* mouse line with *loxP* sites flanking exons 2-10 of the *Tp53* gene was purchased from Jackson’s laboratory (*Trp53^loxP/+^*) (stock# 008462) (Marino S, *et.al*., 2000)

### Generation and establishment of *Snrpb* conditional mutant mouse lines

To develop a conditional knock-out *Snrpb* allele, we used CRISPR/Cas9-mediated homology-directed repair (HDR) strategy to insert *loxP* sequences in intron 1 and intron 3 to flank exons 2 and 3. Candidate efficient guide RNAs were selected based on previous references (Xu et. al., 2015). Microinjection with single-strand DNA template, guide RNAs, and Cas9 mRNA was done. In the first round of microinjection targeting intron 1, a *loxP* sequence was inserted in intron 1 of 25% of the animals born. The insertion was confirmed in two animals (1 male and 1 female) by Sanger sequencing. We generated homozygous animals with *loxP* sequences in intron 1 from those mice. Intron 1 homozygous *loxP* animals were then used for the second round of microinjection to insert *loxP* in intron 3. Sanger sequencing of DNA from a G1 male offspring of a targeted founder from the second round of microinjection and a wild-type CD1 female was used to confirm that both *loxP* sequences in intron 1 and intron 3 were intact. Thereafter, we backcrossed the animals for at least 5 generations to establish the *Snrpb* conditional mutant mouse line and to remove any potential off-target effect from CRISPR editing. All embryo analysis was of embryos on the mix CD1; C57BL/6J genetic background.

### Generation of *Snrpb ^+/-^* mutant embryos

*Snrpb ^+/-^* mutants were generated by crossing *Snrpb^loxp/+^* mice with *β-Actin-Cre^tg/+^* mice.

### Generation neural crest cell-specific *Snrpb ^+/-^* mutants

To generate embryos and animals with neural crest-specific *Snrpb* heterozygosity, *Wnt1-Cre2 ^tg/+^* animals were mated with *Snrpb^loxp/+^* mice. Embryos obtained from these mating were *Snrpb* heterozygous mutant in the neural crest cells and their derivatives, while all other cells were *Snrpb* wild type.

### Genotyping of mice and embryos

Genomic DNA was extracted from mouse tails or yolk sacs by alkaline lysis (Hou, Gupta *et al*., 2017). For *Snrpb*, genotyping was performed to identify the wild-type and conditional allele (with *loxP* sequences) to amplify segments of intron 1 using the following program: 30 sec 95°C, 30 sec 62°C, 30 sec 72°C for 35 cycles followed by an elongation step of 10 minutes at 72°C. This PCR amplified the targeted DNA segment to determine a wild-type (347 bp) and a mutant (387 bp) amplicon. The primers used for the genotyping were: forward-5’ CCCGAGACAGACACAACATAAG 3’, reverse-5’ GCTTTGAAGGTCCCGATGAA 3’. For the commercially available lines, namely R26R, *Wnt1-Cre2*, mT/mG and beta-actin cre genotyping was performed as detailed on Jackson’s laboratory website: protocol# 29915 (*R26R*), #25394 (*Wnt1-Cre2*), #20368 (mT/mG) and ##33618 (beta-actin cre), respectively.

### Collection of embryos

The day of plug was considered embryonic day 0.5 (E0.5). On the day of dissection embryos were dissected from their extraembryonic membranes. Yolk sacs were used for genomic DNA extraction and genotyping. All embryos were assessed for the presence of a heartbeat and somite number was counted between E8.5 and E10.5. Embryos were fixed in 4% paraformaldehyde in PBS at 4°C overnight (unless otherwise stated), washed and kept in PBS at 4°C until use.

### Wholemount in situ hybridization and preparation of embryos for embedding

Fixed embryos were dehydrated using a graded methanol series for wholemounts. Wholemount RNA *in* situ hybridization was performed as previously described (Revil and Jerome-Majewska 2013). For cryo-embedding, fixed embryos were first cryoprotected in 30% sucrose overnight, embedded in Cryomatrix and stored at −80°C until sectioning.

### Cartilage staining of embryos and skeletal preparation of embryos and pups

To investigate cartilage formation, embryos were stained with Alcian Blue (Regeur, 2013). For skeletal staining, the skin was removed from freshly dissected E17.5 embryos and neonatal pups and stained as described by Wallin et al. (1994). We measured the mandible length from the incisor and to the articular surface of the condyloid process. In mutants where the processes of the mandibles were not properly formed, the incisor to the proximal end of the mandible was measured.

### Wholemount X-galactosidase staining of embryos

Embryos were stained with freshly prepared X-gal staining solution overnight at 37°C in the dark as described in Beauchamp *et al*, 2021 Post-staining, embryos were embedded in Cryomatrix and stored at −80°C until sectioning. Image J was used to quantify the lacz stained area.

### PTA staining for CT scan

Embryos were fixed overnight in 4% PFA and then washed with PBS. After a series of dehydration according to the protocol describe previously (Lesciotto *et al*., 2020) embryos were stained in 0.7% Phosphotungstic Acid (PTA). The duration of staining varied depending on the stage of the embryos and pre-scanning was done to confirm complete penetration of the PTA. Once all the structures were visualised in the pre-scan, embryos were rehydrated in a series of methanol and CT scanning was done at 20-micrometer thickness.

### Immunofluorescence and TUNEL assay

Immunofluorescence experiments were performed on 10-micron thick sections according to standard protocols (Zakariyah et al., 2011). The following primary antibodies were used: Phosphohistone H3 (Ser10) [source] (1:200 dilution). Alexa Fluor 568 [source] (1:500 dilutions) secondary antibody was used. For identifying cells undergoing apoptosis, TUNEL assay using Cell Death Detection Kit, TMR red was used (Roche, cat# 12156792910). For quantification of fluorescence signal, particle analysis on Image J was used. For TUNEL and PH3 quantification, at least 4 sections were counted per embryo and per genotype.

### Immunohistochemistry

Embryos were sectioned at 10-micron thickness for immunohistochemistry as previously described (Beauchamp *et al*., 2021; Hou *et al*, 2017). P53 primary antibody (Cell signaling, Cat#2524) was used (1:250 dilution) and the VECTASTAIN® Universal Quick HRP Kit was used as secondary antibody and visualized with DAB.

### RNA isolation for RNA sequencing

RNA extraction was done using Qiagen RNeasy kit following manufacturer’s protocol from samples stored in RNA later (Invitrogen). For RNA isolation at E9.0, heads of two somite-matched embryos from different litters were pooled according to genotype. 3 wild-type and heterozygous pools were used for RNA sequencing analysis.

### qRT-PCR

Total RNA was treated with DNAse (NEB, according to manufacturer’s protocol) and used for reverse transcription with the iScript™ Reverse Transcription Supermix for RT. qRT-PCR was performed using the Advanced universal SYBR® Green Supermix. RT-qPCR experiments were performed in duplicates to ensure technical replicability. Target genes were normalized with the normalization factor as calculated by geNorm software (Vandesompele *et al*. 2002). Three housekeeping genes including B2M, GAPDH, and SDHA were used for generation of the normalization factor as previously reported (Vandesompele *et al*. 2002).

### RNA sequencing Analysis

Sequencing libraries were prepared by McGill Genome Centre (Montreal, Canada), using the TruSeq Stranded Total RNA Sample Preparation Kit (Illumina TS-122-2301, San Diego, California, United States) by depleting ribosomal and fragmented RNA, synthesizing first and second strand cDNA, adenylating the 3’ ends and ligating adaptors, and enriching the adaptor-containing cDNA strands by PCR. The libraries were sequenced using the Illumina NovaSeq 6000 PE100 sequencer, 100 nucleotide paired-end reads, generating between 109 and 230 million reads sample. The sequencing reads were trimmed using CutAdapt (Martin M., 2011) and mapped to the mouse reference genome (mm10) using STAR (Dobin A et al., 2013) aligner (version 2.6.1d), with default parameters, and annotated using the Gencode (Harrow J *et al*., 2006) M2 (version M2, 2013) annotation. Htseq-count (part of the ‘HTSeq’ (Anders S *et al*., 2015) framework, version 0.13. 5) was used for expression quantification.

To perform a differential splicing analysis, rMATS 4.0.2 (Shen S. *et al*., 2014) was used and detected splicing events were filtered by systematically excluding those with a mean of inclusion junction counts (IJC) lower than 5 in either wild-type or heterozygous samples. To identify significant DSE, an absolute inclusion level difference (ILD) cut-off of more than 0.05 was used and a Benjamin-Hochberg multiple testing correction with a False Discovery Rate (FDR) cut-off of less than 0.1 was used. The rationale for relaxing the FDR cut-off here was to obtain a large dataset enriched for alternative splicing events in order to observe general tendencies, such as increased propensity for exon skipping or intron retention in the mutants. To characterize 3’SS sequences, LaBranchoR (Paggi and Bejerano, 2018), a branchpoint (BP) prediction tool based on a deep-learning approach was used, that uses a bidirectional long short-term memory network (LSTM) model, to identify relevant branchpoints upstream DSEs. The BPs and their surrounding area consensus motifs were generated using WebLogo 3.0 (Crooks GE *et al*., 2004).

For differential expression analysis (DEA), we used DESeq2 (Love M.I. *et al*., 2014) package and a list of significant DEG was derived using a FDR cut-off of less than 0.05 with no additional restriction on the absolute log2 fold change (Log2FC) (to allow for detection of even minor expression changes). For KEGG pathway analyses, the combined list of up- and down-regulated genes from DEA was used as input to gProfiler2 (Raudvere U. et al., 2019) package (gost function), and all the detected genes from DEA were used as background.

A differential analysis of transposable element (TE) and lncRNA expression was also carried out, to investigate whether SNRPB deficiency may result in deregulation of the non-coding transcriptome. Those analyses did not uncover any differences in the mutant embryos, and the results are not shown in the manuscript. STAR was used to map the processed reads with modified options:—outFilterMultimapNmax 100 —winAnchorMultimapNmax 100 — outMultimapperOrder Random —alignSJoverhangMin 8 —outFilterMismatchNmax 999 — alignIntronMin 20 —alignIntronMax 1000000 —alignMatesGapMax 1000000, with mouse annotations to guide mapping, coming from the UCSC repeat masker (Gencode M1) and lncRNA (Gencode M1) annotations. The mapped lncRNA and TE reads were respectively quantified with salmon (Patro R. *et al*., 2017) and TElocal (Jin Y *et al*., 2014). Differential lncRNA and TE expression analysis were performed using DESeq2, with the TE and lncRNA read counts being normalized using protein-coding gene expression size factors and differentially expressed lncRNA and TE selected based on a FDR cut-off of less than 0.05 and an absolute Log2FC of greater than 0.5 to increase detection signal.

### Quantification and Statistical analysis

Quantitation was performed using Image J software (NIH). Statistical analyses were conducted using GraphPad Prism 8.0 software (GraphPad Prism, San Diego, CA, USA). Chi-square test or non-parametric Man Whitney t-test analysis was performed using the Prism. Significant p-values are represented as *P<0.05, **P<0.01 and ***P<0.001.

## Legends to Supplementary Figures

**Figure S1: *Snrpb*^ncc+/-^ embryos show craniofacial malformations of varying expressivity from E9.5 onward**. Representative images showing control (*Wnt1-Cre^Tg+^* or *Snrpb^L/+^*) and *Snrpb*^ncc+/-^ (*Snrpb^L/+^; Wnt1-Cre^Tg/+^*) embryos grouped by age and phenotype **A**. At E9.0, all *Snrpb*^ncc+/-^ embryos were morphologically normal and resembled control littermates, they were classified as group 1. **B**. At E9.5, 50% of *Snrpb*^ncc+/-^ embryos were morphologically normal and 50% had hypoplastic forebrain, midbrain and hindbrain and were classified as group 2. **C**. Morphologically normal E10.5, *Snrpb*^ncc+/-^ embryos were assigned to group1, and abnormal embryos that showed reduced frontonasal prominence, small pharyngeal arches, forebrain and midbrain were classified as group 2. **D**. E11.5 *Snrpb*^ncc+/-^ mutants were classified into three groups. Group 1 was morphologically normal; group 2 had hypoplasia of the forebrain, midbrain and hindbrain as wells as reduced maxillary and mandibular prominences; group 3 mutants had severe hypoplasia of the frontonasal, maxillary prominences and the mandibular prominence. **E**-**G**. At E12.5 (**E**), E14.5 (**F**) and E17,5 (**G**), *Snrpb^ncc+/-^* mutants were classified into 4 groups. Group 1 was morphologically normal; group 2 had frontonasal and mandibular clefts; group3 had reduced and abnormal forebrains with clefts in their hypoplastic frontonasal and mandibular prominences; and group 4 mutants did not have a morphologically identifiable forebrain or anterior facial structures. **G**. Group 1 or morphologically normal embryos were not found at E17.5. 1, 2= pharyngeal arch 1 and 2, respectively, fb=forebrain, mb=midbrain, hb=hindbrain, fl=forelimb, E=ear, N=nose, hl=hindlimb.

**Figure S2: E14.5 *Snrpb^ncc+/-^* mutants show hypoplastic and loss of craniofacial cartilage development.** Representative images of E14.5 control (*Snrpnrpb^L/+^*) and *Snrpb^ncc+/-^* embryos stained with Alcian blue **A**. Representative images showing sagittal views of a wild type, control, embryo with normal development of head, nasal and Meckel’s cartilage (white arrowhead); a group 2 *Snrpb^ncc+/-^* group 2 mutant with a shorter and discontinuous Meckel’s cartilage (mc) (red arrowhead); a group 3 mutant with reduced head cartilage (red arrowhead), ectopic cartilage in the maxillary prominence (red arrowhead), and an absent nasal cartilage; a group 4 mutant with absent anterior craniofacial cartilages **B**. Representative images showing ventral views of the head of a wildtype, control, embryo with symmetrical Meckel’s cartilage (white arrowhead) and nasal cartilage; in the group 2 mutant Meckel’s cartilage is truncated and asymmetrical (red arrowhead) and the nasal cartilage is clefted (red star); in group 3 and 4 mutants, ventral cartilages are missing. **C**. and **D**. Higher magnification of the ear (lateral view) and cranial base (ventral view) of control and a group 3 *Snrpb^ncc +/-^* mutant indicating ectopic cartilages found in a subset of mutants (n=4/7), red arrowheads. mc=Meckel’s cartilage, nc=nasal cartilage.

**Figure S3: Craniofacial malformations in newborn *Snrpb^ncc+/-^* mutants**. Representative images of P0 control (*Wnt-Cre^tg/+^*) and *Snrpb^ncc +/-^* pups stained with Alcian blue and Alizarin red **A**. Representative images of ventral views showing normal closed palate and normal bones in the head. The single mutant from group 2 with a bony palate cleft (yellow arrowhead) is shown. This embryo was also missing the tympanic ring and alisphenoid bone; the abnormally shaped basisphenoid bones can also be seen. In the second image, a group 1 mutant with a closed bony palate, but abnormal alisphenoid, basisphenoid and hypoplastic tympanic ring is shown. **B**. Representative images of mandibles of a wild type, control pup shows the normal shape and morphology of the coronoid, angular and condyloid processes. In the *Snrpb^ncc+/-^* mutant from group 2 (mut 1), a symmetrical but abnormal lower jaw is shown. In this same embryo, Meckel’s cartilage was shortened, and the angular processes was missing. In a second mutant from group 2 (mut 2), an asymmetrical lower jaw is shown with Meckel’s cartilage and the angular process missing on one side. **C**. Representative images of lateral views showing normal tympanic ring (arrow) and ossification in the head of a normal embryo. In the representative images of two group 2 *Snrpb^ncc+/-^* mutants the tympanic ring was absent in one and hypoplastic (arrow) in the second. In addition, the bones of the middle ear were abnormal with ectopic bones in the middle ear (white arrowhead) in one mutant and missing in the second. E=ear, Y=eye, BS=basisphenoid bone, AS=alisphenoid bone, PL=palatine, PPPL= palatal process of palatine, PMX=premaxilla, PPPMX=palatal process of premaxilla, PPMX=palatal process of maxilla, Tr=tympanic ring of ear, AP=angular process, CP=coronoid process, CNP=condyloid process, MC=Meckel’s cartilage.

**Figure S4: Micro CT scan of *Snrpb^ncc+/-^* mutants show abnormal development of heart, brain, palate and thymus. A**. Mid-sagittal MicroCT images of a control, (*Snprb^L^/+*) embryo with normal morphological landmarks in the brain and face, and a *Snrpb^ncc+/-^* group 3 mutant with an enlarged lateral ventricle in the brain. The missing nasopharyngeal cavity and nasal septum can also be seen. **B**. and **C.** Transverse MicroCT images of the chest region of an E17.5 control embryo (*Snprb^L^/+*). Panel **C**. show higher magnification of the heart (red box in panel B) from posterior to anterior (left to right of the panel), and the aorticopulmonary septum (white arrowhead) separating the aorta and pulmonary arteries in the control embryo (n=1). **D**. and **E**. Transverse MicroCT images of the chest and heart region of an E17.5 group 2 *Snrpb^ncc+/-^* mutant. **E.** in the higher magnification of the heart (red box in panel D) the aorticopulmonary septum is missing (red arrowhead). **F**. Mid-sagittal view of the same group 2 *Snrpb^ncc+/-^* mutant showing absence of the thymus (red arrowhead) which can be seen in the control embryo (white arrowhead). **G**. Sagittal views of a control and a group 2 E17.5 embryo shows the fused palatal shelves (white arrowheads) in the control, and clefts in the palate and the maxilla in the *Snrpb^ncc+/-^* mutant (red arrowheads). lv=lateral ventricle, Ns =nasal septum, Nc=nasopharyngeal cavity, Pg=pituitary gland, Tg=tongue, Op=oropharynx, LV=left ventricle, RV=right ventricle, Ao=aorta, Pa=pulmonary artery, Bs=basisphenoid bone, Th=thymus, Mx=maxilla

**Figure S5: Neural crest cells number and proliferation are not affected in E9.5 *Snrpb^ncc+/-^* mutants. A**. Representative images of an X-gal stained E9.5 control (*Wnt1-Cre^Tg/+^*) and a group 2 *Snrpb^ncc+/-^* embryos. Fewer X-gal positive cells (blue) were found in the craniofacial region and the pharyngeal arches of the mutant. **B**. and **C**. Representative image of cryosection in a group 2 embryo and quantification of blue cells revealed a non-significant reduction in the percentage of X-gal positive cells in *Snrpb^ncc+/-^*(n=3) embryos when compared to controls (n=3). **D**-**G**. Representative images of DAPI stained E9.5 control (*Snrpb^wt^*) and group 1 *Snrpb^ncc+/-^* embryos carrying the mT/mG reporter. Green-fluorescence marks *Wnt1*-cre-expressing cells. **D.** and **F.** Lower magnification images of whole embryos. **E**. and **G**. Higher magnification of the craniofacial region (boxes in panel D and F) of control and mutant embryos, respectively, showing similar proportion of GFP^+^ cells in the mutant embryo **H**. and **I**. A magnified view of the pharyngeal arch of cryosectioned embryos (n=3) showing presence of GFP^+^ cells (green) in the pharyngeal arches of controls and mutants, respectively. **J** and **K**. Quantification of phosphohistone H3 positive cells in the craniofacial region revealed no significant difference between controls (*Snrpb^wt^*) and mutants at E9.5 or E10.5. At E9.5, n=4 *Snrpb^ncc+/-^* mutants (3 group 1 and 1 group 2). At E10.5 n=5 controls and 5 *Snrpb^ncc+/-^* mutants (1 group 1 and 4 group 2). Error bars indicate standard error of mean (SEM) hb=hindbrain, mb=midbrain, fb=forebrain, 1,2=pharyngeal arch 1 and 2, respectively, pa=pharyngeal arch, nt=neural tube, ncc=neural crest cells, mc=mesenchymal core.

**Figure S6: Increased branch point strength might contribute to aberrant splicing in *Snrpb^ncc+/-^* mutants. A.** and **B.** Presence of a premature termination codons (PTCs) in exons that are more skipped in mutant (**A**) and wild-type embryos (**B**) do not impact exon inclusion level. No significant difference was observed between (more skipped) PTC and Non-PTC exons. Similarly, **C.** and **D.** show no effect of PTCs in exons that are more skipped in mutant (**C**) and wild-type embryos (**D**) on the expression of genes carrying those exons. **E**. and **F**. Splice site weakness assessment (MaxEntScan scores): comparative analysis of the 5 ‘splice site strength (**E**) and 3’ splice site strength (**F**) of introns that are more retained in mutant embryos and wild-type embryos, respectively. No significant difference was found in both analyses between the 2 conditions (Significance cut-offs, p-value <= 0.1[*] or p-value <= 0.05[**]), T-test. **G** and **H**. A 9-mer (3 exonic flanks and 6 intronic flanks around the 5’ss) consensus motif derived from introns that are more retained in mutant embryos (**G**) and wild-type embryos (H). **I** and **J**. A 23-mer (20 intronic flanks and 3 exonic flanks around the 3’ss) consensus motif derived from introns that are more retained in mutant embryos (**I**) and wild-type embryos (**J**). **K**. Highlights a significant difference in the strength of the BP sites (taking into account the LaBranchoR BP scores, 23bp upstream the 3’SS of exons that are more skipped in mutant embryos in comparison of those skipped in the wild-type embryos), T-Test. **L**. A mean comparison test (t-test) of the BP Distance from the 3’SS of the exons that are more skipped in mutant compared to the wild-type embryos. **M**. Highlights a mean comparison test (T-Test) of the LaBranchoR predicted BP Score from the introns that are more retained in mutant embryos in comparison to those more retained in the wild-type embryos, with no significant difference. **N**. A mean comparison test (T-Test) of the BP Distance from the 3’SS of the introns that are more retained in the mutant compared to the wild-type embryos. - Significance cutoffs used: p-value > 0.05 [ns] and p-value <= 0.05[*]. **O** and **P**. A (consensus) motif analysis of the branchpoint site (23bp around the LaBranchoR predicted branchpoint [BP]) from exons that are more skipped in mutant embryos (O) and wild-type embryos (P). **Q** and **R**. A (consensus) motif analysis of the branchpoint sites (23bp around the LaBranchoR predicted BPS) from introns that are more retained in mutant (Q) and wild-type embryos (R). **S** and **T**. show a GC content analysis (G+C nucleotide frequency analysis) of the 23bp sequences, upstream the LaBranchoR predicted BP of exons that are more skipped (S) and more retained (T) in mutant and wild-type embryos. A clear higher GC proportion in the mutant embryos is seen in case of retained intron.

**Figure S7: Knockdown of P53 in the neural crest cells do not rescue craniofacial abnormalities in *Snrpb^ncc+/-^* mutants. A**. and **B.** Representative images of sections of heads of E9.5 control (*Wnt1-Cre^Tg/+^*) and *Snrpb^ncc+/-^* embryos after immunohistochemistry to detect P53 (brown). Increased nuclear P53 was found in mutants (n=3) when compared to controls (n=3). **C**. Similar range of phenotypic abnormalities was found in E17.5 *Snrpb^ncc+/-^* embryos heterozygous for *Trp53* when compared to *Snrpb^ncc+/-^* embryos (shown in Figure 1) **D**. Representative images of skeletal staining of E17.5 control (*Snrpb^L/+^*) and *Snrpb^ncc+/-^; P53^ncc+/-^* embryos showing that P53 heterozygosity in neural crest cells do not rescue craniofacial abnormalities in *Snrpb^ncc+/-^* embryos. **E**. Representative images of E14.5 control (*Snrpb^L/+^*) and *Snrpb^ncc+/-^; P53^ncc-/-^* embryos showing micrognathia and hypoplasia of the head and outer ear in the mutant. **F** Representative images of E14.5 control (*Snrpb^L/+^*) and *Snrpb^ncc+/-^* ; *P53^ncc-/-^* embryos after Alcian blue staining showing a hypoplasia of head and nasal cartilages and a curvy and shorter Meckel’s cartilage in the mutant. **G**. Representative images of E18.5 control and *Snrpb^ncc+/-^; Trp53^ncc-/-^* embryos. *Snrpb^ncc+/-^; Trp53^ncc-/-^* (n=4) had microcephaly, a shorter snout and micrognathia. **H**. and **I**. Representative images of Alcian blue and Alizarin red stained skulls of E18.5 control and E18.5 *Snrpb^ncc+/-^; Trp53^ncc-/-^* embryos. **H.** Cranial base view shows hypoplasia of the nasal bone, cleft palate (arrowhead) and hypoplasia of the tympanic ring in the *Snrpb^ncc+/-^; Trp53^ncc-/-^* mutant. **I.** Top view of the calvaria shows reduced ossification of the frontal bone, reduced ossification, and reduced ossification of the nasal bone in the mutant. **J**. Representative images of the lower jaws of a control and a *Snrpb^ncc+/-^; Trp53^ncc-/-^* mutants showing asymmetrical and abnormal development of the mutant mandible. Arrowhead in control indicates the angular process which is not discernable in the mutant (arrowhead). nt=neural tube, hm=head mesenchyme. E=ear, Y=eye, M=mandible, fl=forelimb, mc=Meckel’s cartilage, nc=nasal cartilage, nb=nasal bone, tr=tympanic ring, pl=palate.

