## Supplemental figures and table for "*Snrpb*, the CCMS gene, is required in neural crest cells for proper splicing of genes essential for craniofacial morphogenesis"

FIGURE S1

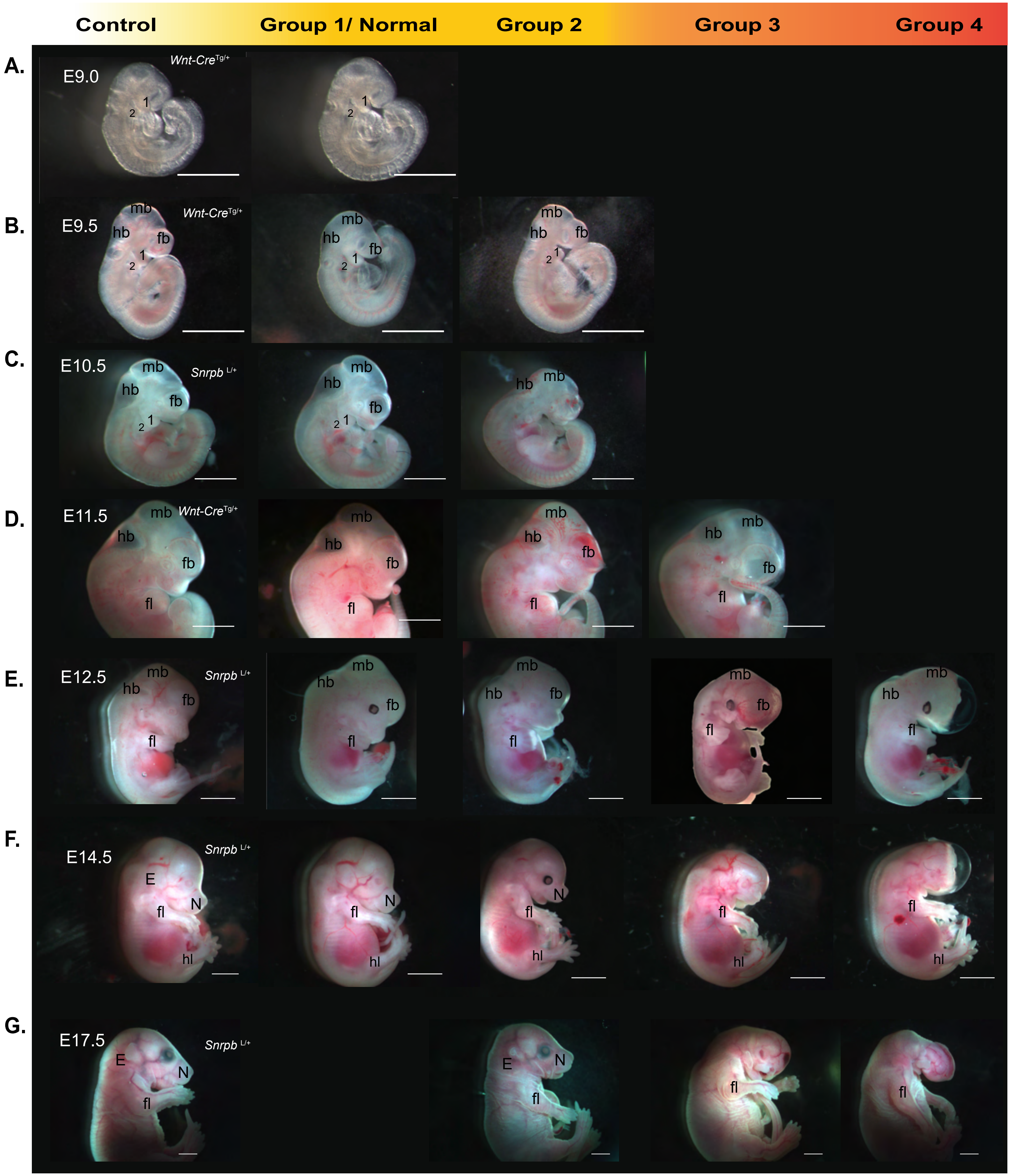
Control
Group 1/ Normal
Group 2
Group 3
Group 4
A.
E9.0
Wnt-Cre Tg/+
B.
E9.5
Wnt-Cre Tg/+
mb
hb
fb
C.
E10.5
Snrpb L/+
D.
E11.5
Wnt-Cre Tg/+
fl
E.
E12.5
Snrpb L/+
F.
E14.5
Snrpb L/+
E
N
hl
G.
E17.5
Snrpb L/+

FIGURE S2

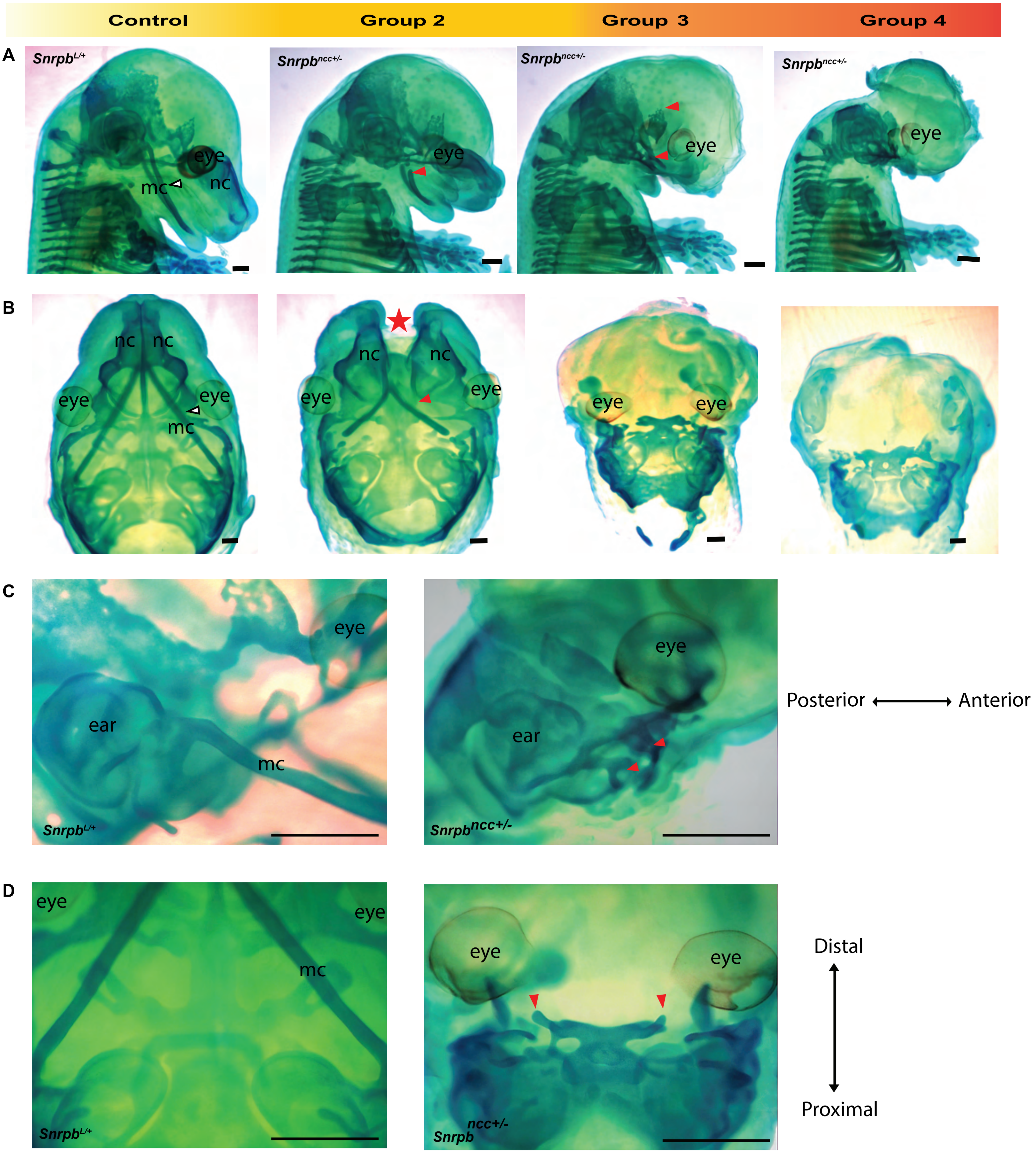

Control
Group 2
Group 3
Group 4
A
Snrpb^L/+
Snrpb^ncc+/-
Snrpb^ncc+/-
Snrpb^ncc+/-
eye
mc
nc
eye
eye
eye
B
nc nc
eye
eye
mc
nc
nc
eye
eye
eye
eye
C
eye
ear
mc
Snrpb^L/+
eye
ear
Snrpb^ncc+/-
Posterior
Anterior
D
eye
eye
mc
Snrpb^L/+
eye
eye
Snrpb ncc+/-
Distal
Proximal

### FIGURE S3

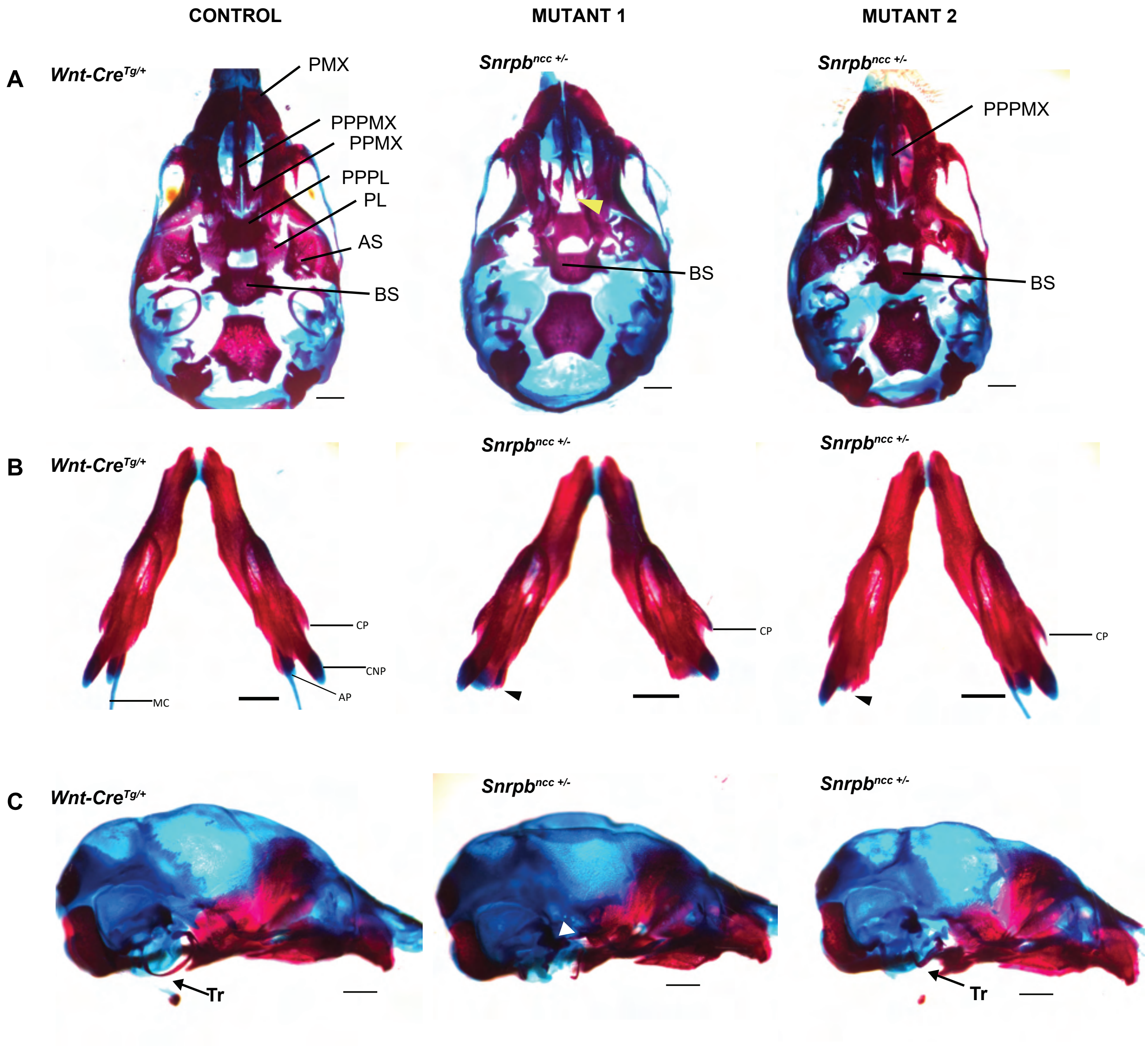

CONTROL
MUTANT 1
MUTANT 2
A
Wnt-Cre$^{Tg/+}$
Snrpb$^{ncc +/-}$
Snrpb$^{ncc +/-}$
PMX
PPPMX
PPMX
PPPL
PL
AS
BS
BS
PPPMX
BS
B
Wnt-Cre$^{Tg/+}$
Snrpb$^{ncc +/-}$
Snrpb$^{ncc +/-}$
CP
CNP
AP
MC
CP
CP
C
Wnt-Cre$^{Tg/+}$
Snrpb$^{ncc +/-}$
Snrpb$^{ncc +/-}$
Tr
Tr

FIGURE S4

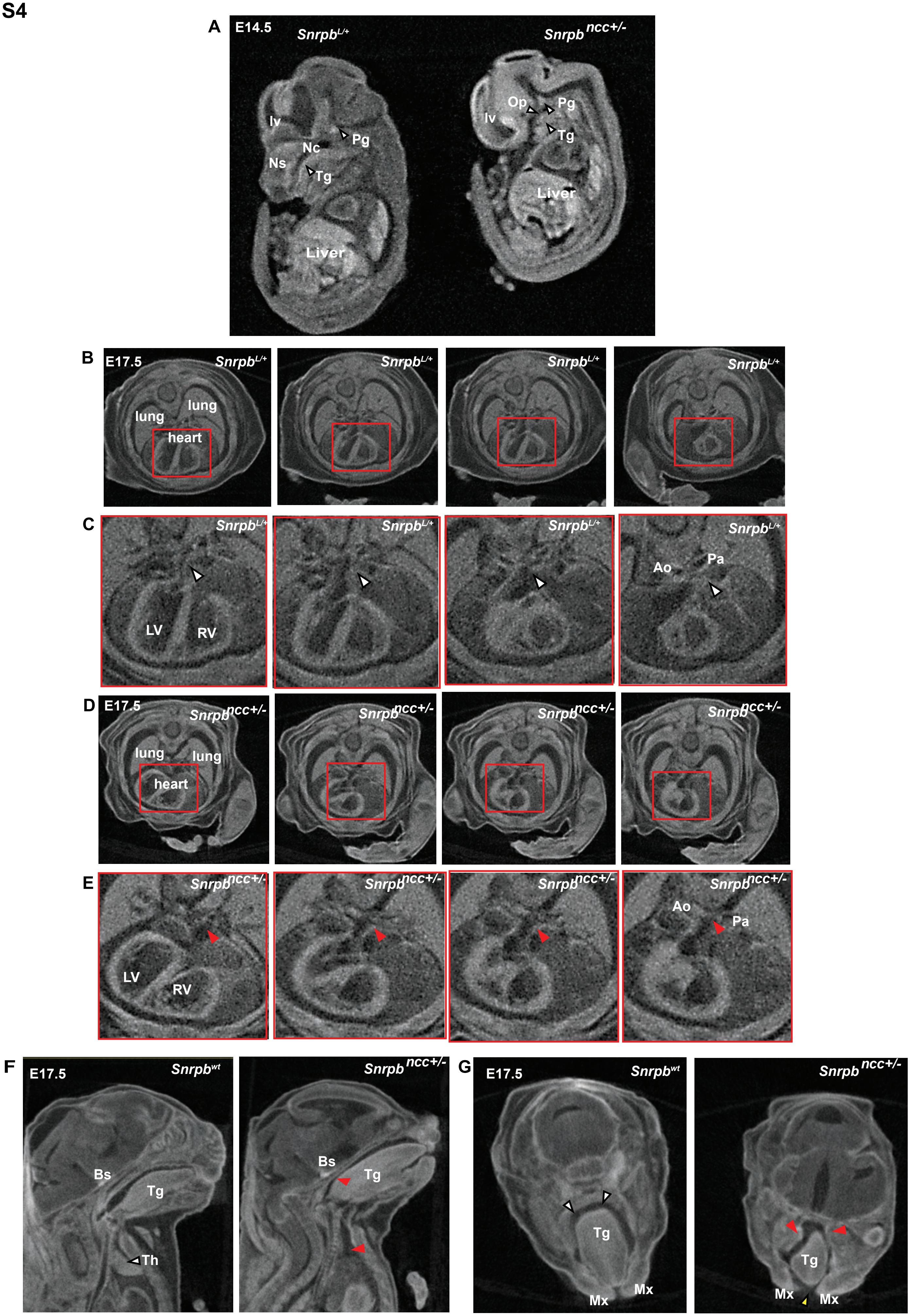
A
E14.5
Snrpb^L/+
Snrpb ncc+/-
lv
Nc
Ns
Pg
Tg
Liver
Op
B
E17.5
lung
heart
C
LV
RV
Ao
Pa
D
E
F
Bs
Tg
Th
G
Snrpb^wt
Mx

### FIGURE S5

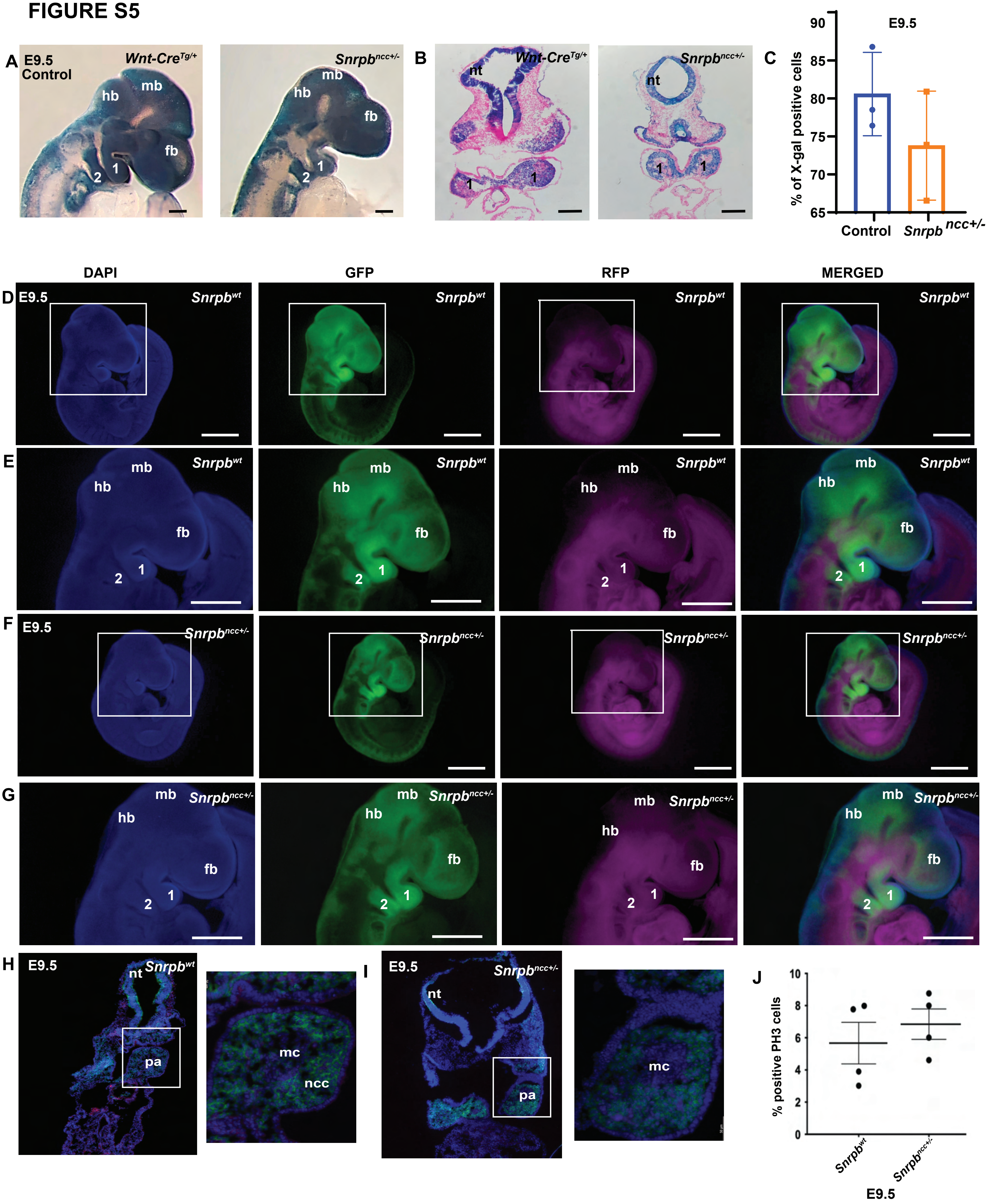
A
E9.5
Control
Wnt-Cre^Tg/+
Snrpb^ncc+/-
hb
mb
fb
1
2
B
nt
C
E9.5
% of X-gal positive cells
90
85
80
75
70
65
Control
Snrpb ncc+/-
DAPI
GFP
RFP
MERGED
D
E
F
G
Snrpb^wt
H
I
pa
mc
ncc
J
% positive PH3 cells
10
8
6
4
2
0

### FIGURE S6

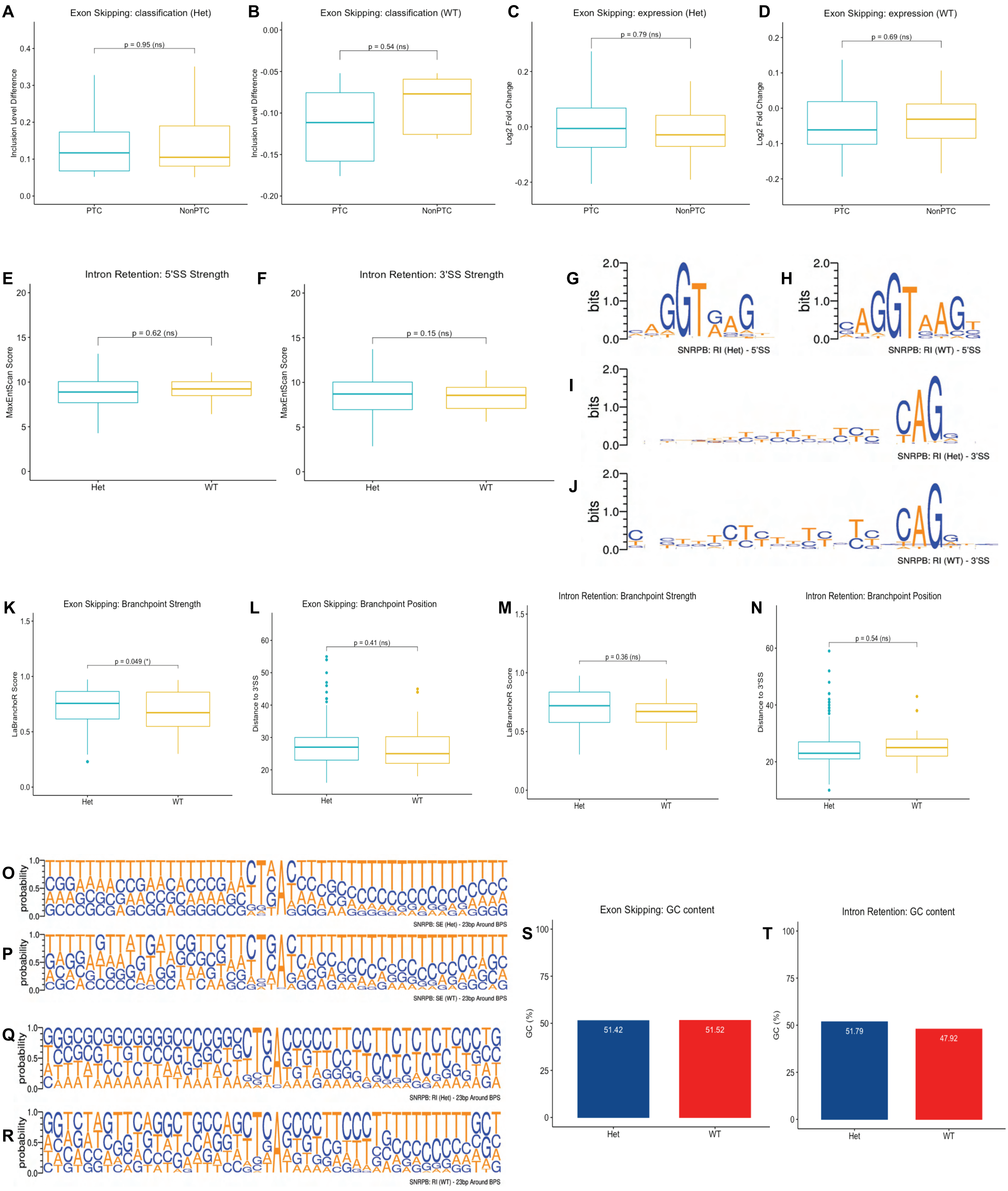
A
Exon Skipping: classification (Het)
Inclusion Level Difference
p = 0.95 (ns)
PTC
NonPTC
B
Exon Skipping: classification (WT)
Inclusion Level Difference
p = 0.54 (ns)
PTC
NonPTC
C
Exon Skipping: expression (Het)
Log2 Fold Change
p = 0.79 (ns)
PTC
NonPTC
D
Exon Skipping: expression (WT)
Log2 Fold Change
p = 0.69 (ns)
PTC
NonPTC
E
Intron Retention: 5'SS Strength
MaxEntScan Score
p = 0.62 (ns)
Het
WT
F
Intron Retention: 3'SS Strength
MaxEntScan Score
p = 0.15 (ns)
Het
WT
G
bits
SNRPB: RI (Het) - 5'SS
H
bits
SNRPB: RI (WT) - 5'SS
I
bits
SNRPB: RI (Het) - 3'SS
J
bits
SNRPB: RI (WT) - 3'SS
K
Exon Skipping: Branchpoint Strength
LaBranchoR Score
p = 0.049 (*)
Het
WT
L
Exon Skipping: Branchpoint Position
Distance to 3'SS
p = 0.41 (ns)
Het
WT
M
Intron Retention: Branchpoint Strength
LaBranchoR Score
p = 0.36 (ns)
Het
WT
N
Intron Retention: Branchpoint Position
Distance to 3'SS
p = 0.54 (ns)
Het
WT
O
probability
SNRPB: SE (Het) - 23bp Around BPS
P
probability
SNRPB: SE (WT) - 23bp Around BPS
Q
probability
SNRPB: RI (Het) - 23bp Around BPS
R
probability
SNRPB: RI (WT) - 23bp Around BPS
S
Exon Skipping: GC content
GC (%)
51.42
51.52
Het
WT
T
Intron Retention: GC content
GC (%)
51.79
47.92
Het
WT

FIGURE S7

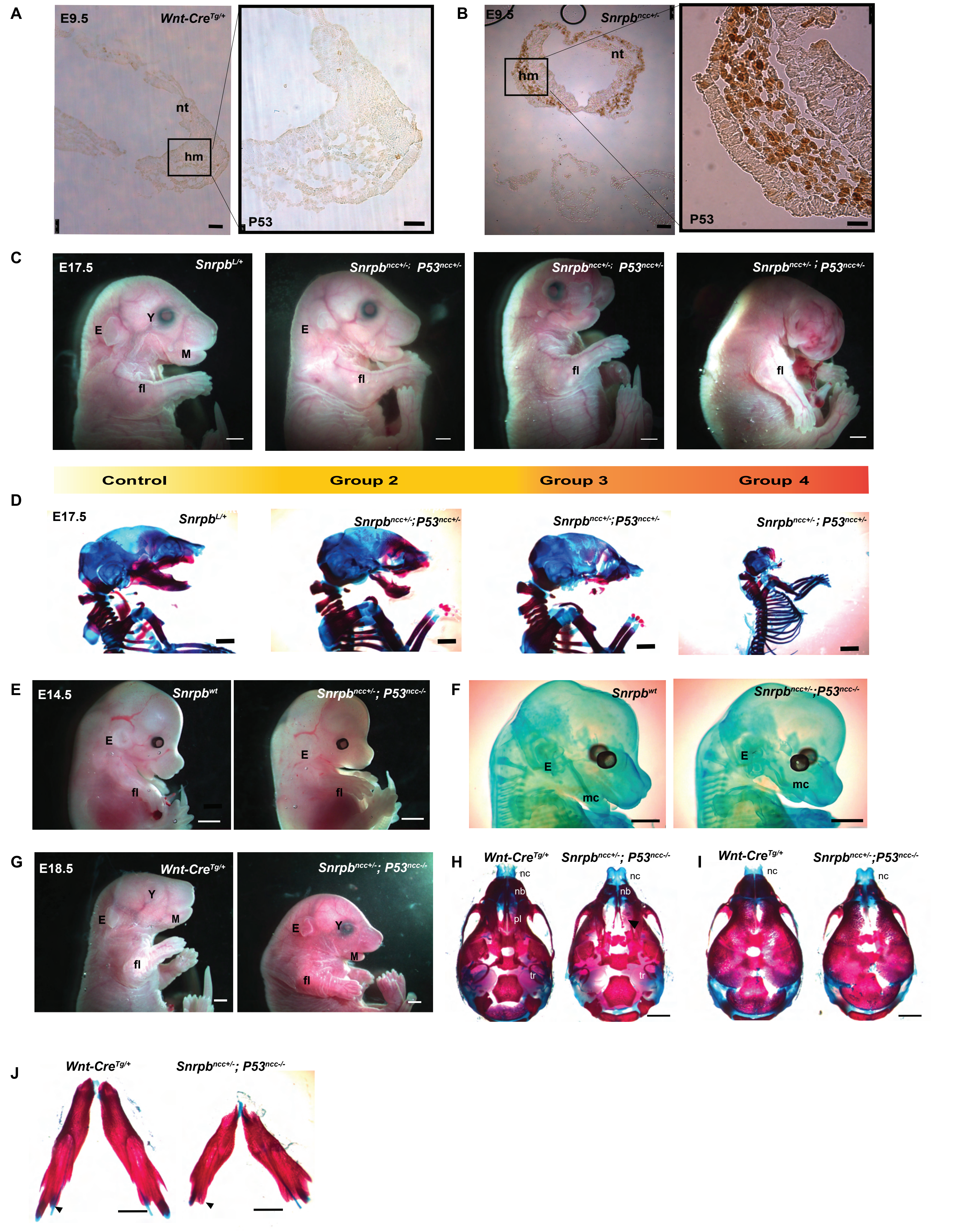

A
E9.5
Wnt-Cre Tg/+
nt
hm
P53
B
E9.5
Snrpb ncc+/-
nt
hm
P53
C
E17.5
Snrpb L/+
Y
E
M
fl
Snrpb ncc+/-; P53 ncc+/-
E
fl
Snrpb ncc+/-; P53 ncc+/-
fl
Snrpb ncc+/- ; P53 ncc+/-
fl
Control
Group 2
Group 3
Group 4
D
E17.5
Snrpb L/+
Snrpb ncc+/-;P53 ncc+/-
Snrpb ncc+/-;P53 ncc+/-
Snrpb ncc+/- ; P53 ncc+/-
E
E14.5
Snrpb wt
E
fl
Snrpb ncc+/-; P53 ncc-/-
E
fl
F
Snrpb wt
E
mc
Snrpb ncc+/-;P53 ncc-/-
E
mc
G
E18.5
Wnt-Cre Tg/+
Y
E
M
fl
Snrpb ncc+/-; P53 ncc-/-
E
Y
M
fl
H
Wnt-Cre Tg/+
nc
nb
pl
tr
Snrpb ncc+/-; P53 ncc-/-
nc
nb
tr
I
Wnt-Cre Tg/+
nc
Snrpb ncc+/-;P53 ncc-/-
nc
J
Wnt-Cre Tg/+
Snrpb ncc+/-; P53 ncc-/-

**Table S1. Increased SE of craniofacial developmental genes in *Snrpb* $^{ncc+/-}$ mutants.** Transcripts required for normal head/craniofacial development with a significant increase in skipped exons are not all predicted to result in PTC in *Snrpb*$^{ncc+/-}$.

| Gene name | Skipped Exon | PTC | Phenotype | Constitutive exon |
|---|---|---|---|---|
| 1. *Smad2* | Exon 3 | No | Mandible hypoplasia (Nomura and Li, 1998) | No |
| 2. *Loxl3* | Exon 2 | No | Cleft palate, short and bent mandible (Zhang *et al.*, 2015) | No |
| 3. *Ror2* | Exon 8 | No | Midface hypoplasia, truncated Meckel's, middle ear defect (Schwabe, G.C. *et al.*, 2004) | Yes |
| 4. *Nisch* | Exon 6 | No | Short snout (Crompton, M. *et al.*, 2017) | Yes |
| 5. *Pou2f1* | Exon 4 | No | Abnormal nasal placode development when removed with *Sox2* (Donner, A.L. *et al.*, 2006) | Yes |
| 6. *Rgl1* | Exon 3 | No | Abnormal frontal bone, short snout, abnormal maxilla and mandibular morphology (Mouse Genome Informatics and the International Mouse Phenotyping Consortium, 2014) | Yes |
| 7. *Frem1* | Exon 32 | No | Midface hypoplasia, asymmetry, short snout (Vissers, L.E. *et al.*, 2011) | Yes |
| 8. *Smc3* | Exon 5 | No | Upturned snout (White, J.K. *et al.*, 2013) | Yes |
| 9. *Pdpk1* | Exon 3 | Yes | Abnormalities in the head, nasal cartilage (Lawlor, M.A. *et al.*, 2002) | Yes |
| 10. *Rere* (*Atr2*) | Exon 4 | Yes | Small pharyngeal arch (Zoltewicz, J.S. *et al.*, 2004) | Yes |
| 11. *Mcph1* | Exon 13 | Yes | Microcephaly (Gruber, R. *et al.*, 2011) | Yes |
| 12. *Nf1* | Exon 56 | Yes | Head hyperplasia, aorticopulmonary septal defect, heart defects (Brannan, C.I. *et al.*, 1994) | Yes |
| 13. *Dyrk2* | Exon 2 | Yes | Cleft palate (Yoshida, S. *et al.*, 2020) | No |
